# Reliability of an interneuron response depends on an integrated sensory state

**DOI:** 10.1101/726083

**Authors:** May Dobosiewicz, Cornelia I. Bargmann

## Abstract

The central nervous system transforms sensory information into representations that are salient to the animal. Here we define the logic of this transformation in a *Caenorhabditis elegans* integrating interneuron. AIA interneurons receive input from multiple chemosensory neurons that detect attractive odors. We show that reliable AIA responses require the coincidence of two sensory inputs: activation of AWA olfactory neurons that are activated by attractive odors, and inhibition of one or more chemosensory neurons that are inhibited by attractive odors. AWA activates AIA through an electrical synapse, while the disinhibitory pathway acts through glutamatergic chemical synapses. The resulting AIA interneuron responses have uniform magnitude and dynamics, suggesting that AIA activation is a stereotyped response to an integrated stimulus. Our results indicate that AIA interneurons combine sensory information using AND-gate logic, requiring coordinated activity from multiple chemosensory neurons. We propose that AIA encodes positive odor valence based on an integrated sensory state.

## Introduction

Sensory environments are complex, and can include multiple signals within and across sensory modalities. Individual stimuli and the combinations in which they occur may differentially signal the presence of beneficial, harmful, or neutral conditions. Accordingly, integration across sensory inputs is an essential function of the nervous system, and the logic of integration is an area of active investigation.

The compact *Caenorhabditis elegans* nervous system, with 302 neurons and a complete neuronal wiring diagram, is well-suited for studying sensory representation and integration (White et al., 1986; Cook et al., 2019). *C. elegans* uses volatile and water-soluble cues to detect its food and other relevant stimuli (Bargmann, 2006). Its highly developed chemosensory system senses a large number of attractive and repulsive compounds using over a thousand different G-protein-coupled chemoreceptors, many of which are expressed by eleven pairs of chemosensory neurons in the amphid sensory organ (Robertson and Thomas, 2006). Specific subsets of chemosensory neurons have reproducible functions in chemotaxis and avoidance of chemical stimuli, spontaneous locomotion, physiology, development, and lifespan (Bargmann, 2006). AWA and AWC sensory neurons, for example, drive chemotaxis toward a variety of attractive odors, whereas AWB sensory neurons drive avoidance of certain repellents (Bargmann et al., 1993; Troemel et al., 1997). Notably, sensory neurons may be either activated or inhibited by odors. For example, AWA is activated by the attractive odor diacetyl (Shinkai et al., 2011; Larsch et al., 2013), whereas AWC is inhibited by another attractive odor, isoamyl alcohol (Chalasani et al., 2007).

At the first layer of neuronal integration, the sensory neurons form abundant chemical and electrical synapses onto a half-dozen pairs of interneurons (White et al., 1986). The interneurons regulate behaviors such as reversal frequency, speed, and head turning during thermotaxis, chemotaxis, and spontaneous locomotion (Iino and Yoshida, 2009; Tsalik and Hobert, 2003; Gray et al., 2005; Hendricks et al., 2012). Interneuron responses are variable and complex, and can incorporate feedback from motor systems as well as sensory input (Gordus et al., 2015; Li et al., 2014; Hendricks et al., 2012; Kaplan et al., 2018). Synaptic mechanisms linking interneurons to locomotor states have been defined, but the computations that interneurons perform to integrate multiple sensory inputs are unknown.

The AIA interneuron pair receives chemical or electrical synapses from all eleven pairs of amphid chemosensory neurons, suggesting an integrative function (White et al., 1986) (Table S1). At a behavioral level, AIA is implicated in the suppression of reversal behavior upon odor addition (Larsch et al., 2015), integration of attractive and repulsive stimuli (Shinkai et al., 2011), olfactory desensitization at short and long timescales (Chalasani et al., 2010; Cho et al., 2016), and regulation of spontaneous reversals (López-Cruz et al., 2019). Functional calcium imaging has demonstrated that AIA can be activated by the attractive odors diacetyl and isoamyl alcohol (Larsch et al., 2013; Chalasani et al., 2010), which are primarily sensed by AWA and AWC neurons, respectively (Bargmann et al., 1993). In both cases, AIA activity rises in the presence of an attractive food-related odor.

Here, we use an optogenetic approach to isolate the connections between individual sensory neurons and AIA. We find that AIA uses AND-gate logic to integrate sensory information. AIA is reliably activated only by coordinated sensory information from multiple neurons, and this activation is mediated by both chemical and electrical synapses. Our results suggest that AIA represents a positive valence that is integrated across sensory inputs.

## Results

### Optogenetic activation of AWA sensory neurons elicits unreliable AIA calcium responses

AWA sensory neurons expressing genetically-encoded calcium indicators such as GCaMP respond to the bacterial odorant diacetyl with fluorescence increases indicating depolarization (Shinkai et al., 2011; Larsch et al., 2013; Zaslaver et al., 2015; Hale et al., 2016; Larsch et al., 2015; Liu et al., 2018). AWA calcium responses are concentration-dependent, with stronger and more rapid responses to 115 nM diacetyl than to 11.5 nM diacetyl, and a rapid rise followed by desensitization within 10 s at 1.15 µM diacetyl (Larsch et al., 2015) (Figures 1A and 1B). Diacetyl also elicits calcium transients in the AIA interneurons, with desensitization at high diacetyl concentrations (Larsch et al., 2015) (Figure 1C). AIA calcium transients are diminished in animals with defects in AWA development or in the AWA diacetyl receptor ODR-10 (Larsch et al., 2015).

**Figure 1.**
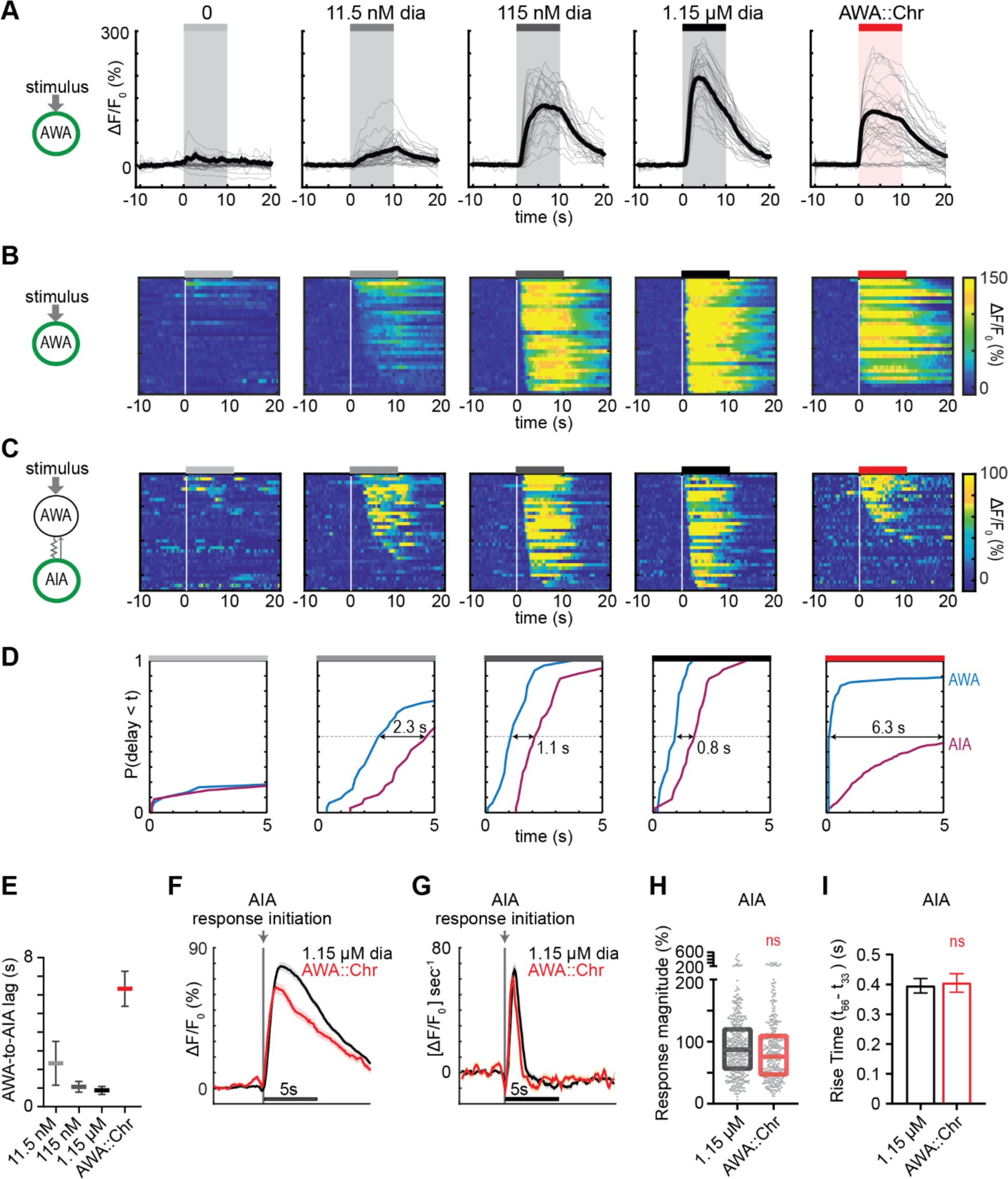
Optogenetic activation of AWA sensory neurons elicits unreliable AIA calcium responses. **(A)** Individual AWA GCaMP2.2b calcium responses to 10 s pulses of increasing concentrations of diacetyl and to optogenetic stimulation. Bold lines indicate mean. Responses to optogenetic stimulation were randomly downsampled to 40 traces from a complete set of 268 traces to match the number of odor traces and enhance visibility. In all schematic diagrams, calcium was monitored in the neuron indicated in green. **(B)** Heat map of AWA calcium responses from (A); responses to optogenetic stimulation were downsampled to 32 traces for visibility; see Figure 1 – figure supplement 1E for complete data. Each heat map row represents a calcium trace to a single stimulus pulse; each animal received two stimulus pulses. Traces are ordered by response latency. **(C)** Heat map of AIA GCaMP5A calcium responses to 10 s pulses of diacetyl or to AWA::Chrimson stimulation. Responses to AWA::Chrimson stimulation were downsampled to 34 traces for visibility; see Figure S1E for complete data. Resistor symbol in cartoon represents a predicted electrical synapse; thin arrow represents chemical synapse. **(D)** Cumulative response time profiles of AWA and AIA responses representing response latencies and probability, without downsampling. Only first 5 s of stimulation are shown. Arrows indicate the delay between the time at which 50% of AWA neurons responded versus the time at which 50% of AIA neurons responded. **(E)** Delay between the time at which 50% of AWA versus 50% of AIA neurons responded to various stimuli, as shown in (D). Delay was greatest to AWA::Chrimson stimulation, despite the short latencies of AWA responses. Bars are mean ± SEM. **(F and G)** Mean AIA responses (F) and corresponding time derivatives (G) to 1.15 µM diacetyl and AWA::Chrimson stimulation that resulted in activation within 5 s of stimulus, aligned to the frame at which activation began and combined over all experiments. Shading indicates ± SEM. 1.15 µM diacetyl: n = 390; AWA::Chrimson: n = 260. **(H)** Magnitude of responses from (F). Boxes show median and interquartile range. (I) Rise time of responses from (F). Bars indicate SEM. For (H) and (I), ns signifies lack of significance of an unpaired t-test. See Table S3 for sample sizes and test details.

To examine AWA-to-AIA synaptic communication in detail, we directly depolarized AWA using the light-activated channel Chrimson expressed under an AWA-selective promoter and recorded GCaMP responses in either AWA or AIA (Klapoetke et al., 2014) (Figure 1A, Figure 1 – figure supplement 1A). Light-activated calcium transients were elicited within one second in Chrimson-expressing AWA neurons and desensitized only slightly over ten seconds. This activation required pre-exposure to retinal and expression of the Chrimson transgene (Figure 1 – figure supplement 1B-C), and was comparable in magnitude to the response of AWA at or above 115 nM diacetyl.

To our surprise, AIA responses to AWA::Chrimson stimulation were significantly smaller than AIA responses to any concentration of diacetyl (Figure 1 – figure supplement 1D). To understand the discrepancy between optogenetic stimulation and odor, we examined the dynamics of individual calcium responses instead of averaged traces (Figures 1B, 1C, and Figure 1 – figure supplement 1E). The GCaMP fluorescence baseline in AIA was stable, with little spontaneous activity, and nearly all AIA neurons responded to the higher odor concentrations (115 nM and 1.15 µM diacetyl) with an average delay of ∼1 second relative to AWA activation (Figure 1B-1E). At the lowest tested diacetyl concentration (11.5 nM) AIA responded with a lower probability (Figures 1C and 1D), and a delay of ∼2 seconds relative to AWA responses (Figure 1E). Thus AIA responses to diacetyl are strongly coupled to odor and dose-dependent. These AIA properties differ from those of AIB interneurons, which have spontaneous calcium fluctuations and variable sensory responses at all odor concentrations due to a strong effect of downstream motor state (Gordus et al., 2015; Kato et al., 2015).

At an individual trial level, AWA::Chrimson stimulation elicited AIA responses with a low probability and a delay, and these responses were much less robust than AIA responses elicited by diacetyl at a comparable level of AWA activation (Figures 1C and 1D). AWA::Chrimson elicited calcium increases in 85% of AWA sensory neurons within one second of optogenetic stimulation, resembling 115 nM diacetyl, but only 56% of the AIA interneurons were activated. Moreover, the AIA calcium transients were delayed by >6 seconds on average relative to the AWA response (Figures 1B-1E). Thus AIA calcium responses to AWA::Chrimson were unreliable.

Control experiments indicated that these differences were robust to transgenes or stimulus protocols. AWA::Chrimson animals had normal AIA responses to diacetyl, before or after light stimulation (Figure 1 – figure supplement 1F), and the AIA response latency was not correlated with GCaMP fluorescence levels or AWA::Chrimson transgene expression levels (Figure 1 – figure supplement 1G-H). Animals were routinely subjected to two stimulus pulses of light or odor; these responses were slightly biased toward the first stimulus but largely independent, indicating that reliability is primarily a trial-to-trial property and not due to variation between animals (Figure 1 – figure supplement 1I-Q).

Close examination of the AIA calcium signals revealed that the response dynamics to diacetyl or AWA::Chrimson stimulation were indistinguishable; only their probability and latency were changed. When aligned to the beginning of an AIA calcium response, positive AIA trials had similar magnitudes, rise times, and decay, whether elicited by optogenetic stimulation or by diacetyl (Figures 1F-1I).

### Gap junctions mediate AWA-to-AIA communication

AWA cell fate mutants (*odr-7*) and AWA diacetyl receptor mutants (*odr-10*) have diminished AIA interneuron responses to diacetyl (Larsch et al., 2015; Sengupta et al., 1994, Sengupta et al., 1996). At the individual trial level, AIA interneuron responses to high (1.15 µM) diacetyl were decreased in magnitude and less reliable in AWA-defective mutants than in wild type (Figure 2A, Figure 2 – figure supplement 1A-B). These results indicate that AWA is necessary for strong and reliable AIA interneuron responses to diacetyl, although the optogenetic experiments indicate that AWA activation is not sufficient for reliable AIA responses.

**Figure 2.**
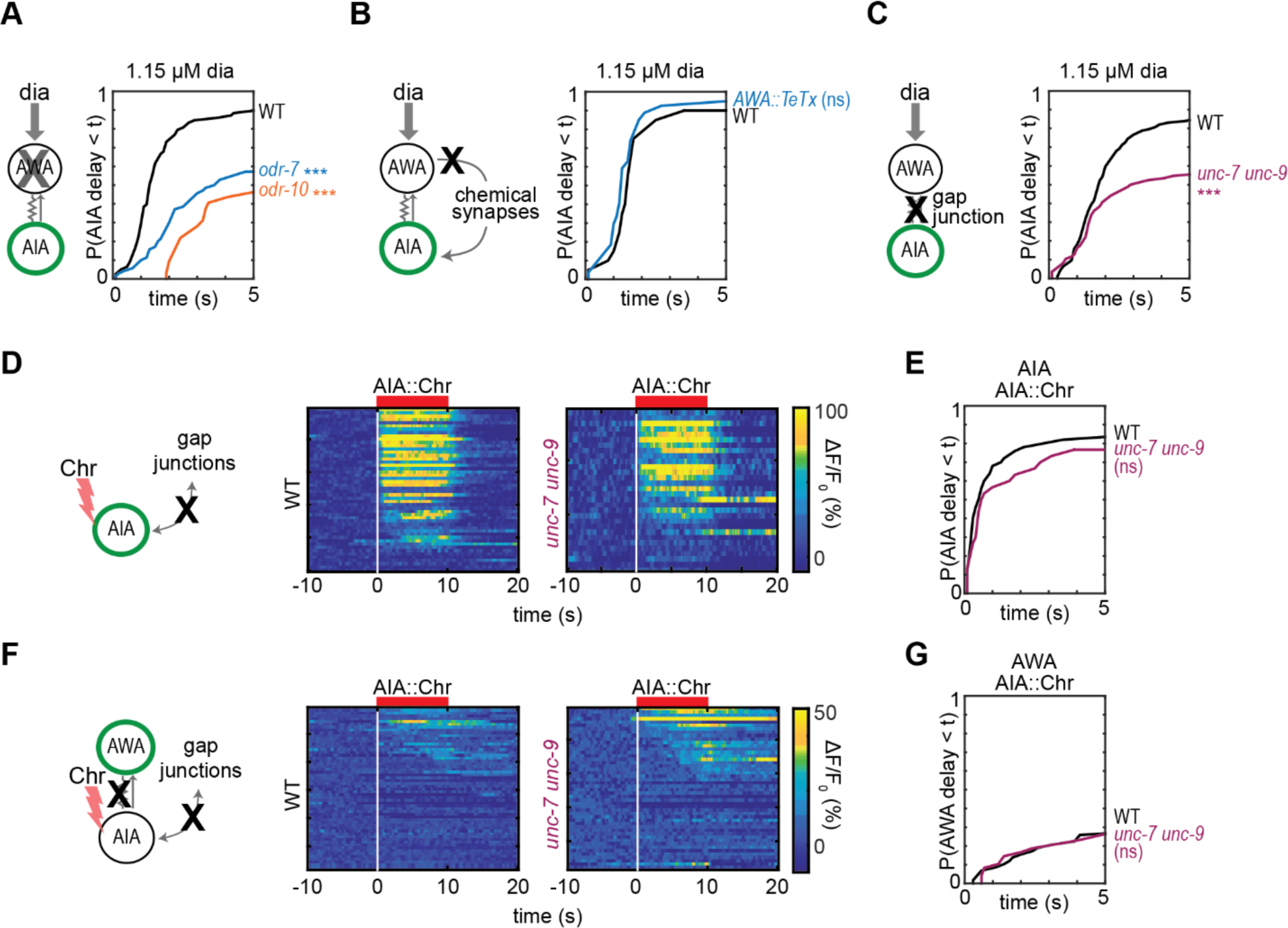
Gap junctions mediate AWA-to-AIA communication. **(A - C)** Cumulative response time profiles of AIA responses to 1.15 µM diacetyl in WT versus *odr-7* animals (AWA cell fate mutants) (A), *odr-10* animals (AWA diacetyl receptor mutants) (A), animals expressing a transgene encoding Tetanus Toxin Light Chain A (TeTx) in AWA (B), and *unc-7 unc-9* animals (innexin mutants) (C). **(D and F)** AIA (D) and AWA (F) responses to 10 s pulses of AIA::Chrimson stimulation in WT and *unc-7 unc-9* animals; one row per calcium trace. Note that scale bar in (F) differs from scale bar in Figure 1B. **(E)** Cumulative response time profiles of AIA responses shown in (D). **(G)** Cumulative response time profiles of AWA responses shown in (F). Asterisks refer to Kolmogorov-Smirnov test significance versus WT over full 10 s stimulus pulse. ns: not significant; ***: p<0.001. See Table S2 for sample sizes and test details.

The *C. elegans* wiring diagram predicts the existence of gap junctions between AWA and AIA neurons, and a recent reanalysis additionally predicts a chemical synapse from AWA to AIA (White et al., 1986; Cook et al., 2019). To assess the importance of chemical synapses, we inhibited AWA synaptic vesicle release by expressing the tetanus toxin light chain in AWA (*AWA::TeTx*), and examined AIA responses to 1.15 µM diacetyl and AWA::Chrimson stimulation. *AWA::TeTx* animals and wild type animals had indistinguishable AIA responses to odor or optogenetic stimuli (Figure 2B, Figure 2 – figure supplement 1C-E), indicating that AWA does not require chemical synapses to activate AIA.

We next asked whether AWA-to-AIA communication requires gap junctions, which are composed of innexin proteins in invertebrates. AWA and AIA express two innexin genes that contribute to many gap junctions, *unc-7* and *unc-9,* among other innexins (Cao et al., 2017; Bhattacharya et al., 2019). AIA responses to diacetyl were less reliable in *unc-7 unc-9* innexin double mutants than in wild type (Figures 2C and S2G), and AIA response magnitudes to diacetyl were decreased (Figure 2 – figure supplement 1F-H), in each case recapitulating the defects observed in AWA cell fate and receptor mutants (*odr-7, odr-10*; Figure 2A, Figure 2 – figure supplement 1A-B). AIA responses to AWA::Chrimson stimulation were also less reliable in *unc-7 unc-9* innexin double mutants, but response magnitudes were unchanged (Figure 2 – figure supplement 1I-J). Together, these results indicate that AWA signals to AIA primarily via gap junctions.

Gap junctions can transmit information symmetrically or asymmetrically between neurons based on properties including voltage-dependent rectification, differential subunit expression between cells, and differential phosphorylation (Goodenough and Paul, 2009). To test whether the predicted electrical synapse between AWA and AIA is bidirectional, we optogenetically stimulated AIA interneurons with a Chrimson transgene and recorded the resulting responses in AWA sensory neurons. AIA responded rapidly and robustly to direct optogenetic stimulation (Figures 2D and 2E), but AWA responded infrequently, with only small-magnitude calcium responses, to AIA stimulation (Figures 2F and 2G). The AWA response to AIA stimulation was unchanged in *unc-7 unc-9* innexin double mutants, and slightly increased in synaptic transmission mutants, suggesting that multiple synapse types contribute to weak AIA-to-AWA communication (Figures 2D-2G, Figure 2 – figure supplement 1K-N). Together, these results suggest that AWA-to-AIA gap junctions primarily mediate anterograde information flow from sensory neurons to interneurons.

### Chemical synapses inhibit AIA

The rapid AIA response to direct optogenetic stimulation indicates that its delayed response to sensory stimuli is not caused by slow intrinsic calcium dynamics, but by other circuit elements. Diacetyl can activate AIA, albeit less reliably, in mutants that lack AWA function or *unc-7* and *unc-9* innexins. The residual AIA diacetyl response indicates that additional diacetyl-sensing neurons communicate with AIA, most likely through chemical synapses. Indeed, AIA receives chemical synapses from many chemosensory neurons (Table S1).

We asked how chemical synapses impact AIA responses by examining mutants in *unc-13* and *unc-18*, both of which are deficient in synaptic vesicle release (Richmond, 2007). Unexpectedly, AIA responses to diacetyl were faster and more reliable in animals with defective chemical synapses (Figure 3A, Figure 3 – figure supplement 1A). This effect was most evident at low diacetyl concentrations, where the synaptic mutants substantially decreased the latency of the AIA response (Figure 3A). Inactivation of dense core vesicle release with an *unc-31* mutation did not alter AIA reliability, demonstrating specificity of the effect (Richmond, 2007) (Figure 3 – figure supplement 1J). Chemical synapses are thus net inhibitory onto AIA.

**Figure 3.**
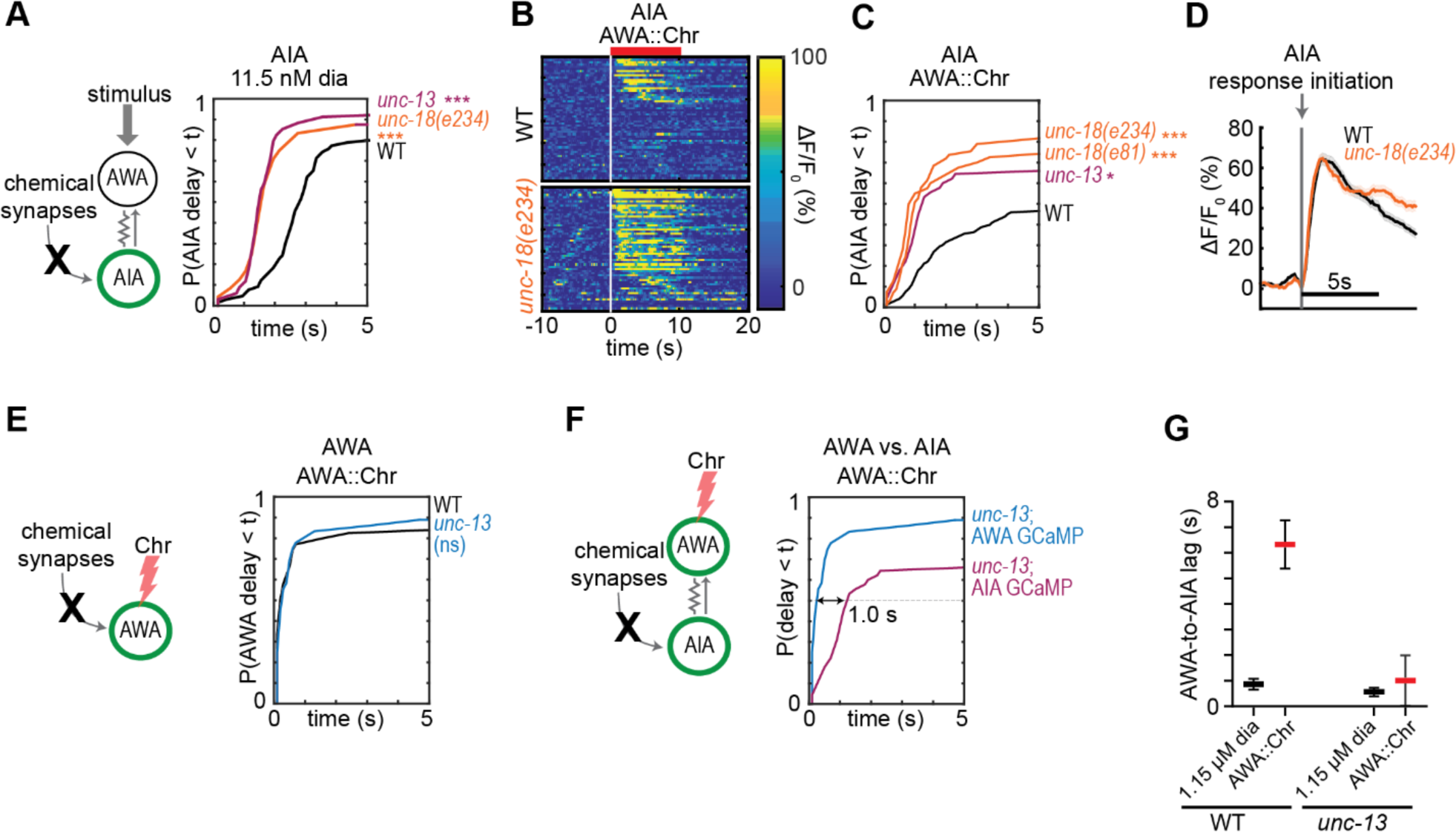
Chemical synapses inhibit AIA. **(A)** Cumulative response time profiles of AIA responses to 11.5 nM diacetyl in WT versus *unc-13(e51)* and *unc-18(e234)* animals (synaptic transmission mutants). **(B)** Heat maps of AIA responses to AWA::Chrimson stimulation in WT and *unc-18(e234)* animals. WT data were randomly downsampled to 57 traces for visibility; data shown for single experiment block. See Figure 3 – figure supplement 1B for pooled data from all experiments. **(C)** Cumulative response time profiles of AIA responses to AWA::Chrimson stimulation in WT versus *unc-13, unc-18(e234),* and *unc-18(e81)* animals. **(D)** Mean AIA responses to AWA::Chrimson stimulation in WT and *unc-18(e234)* animals that resulted in activation within 5 s of stimulus, aligned to the frame at which activation began, combined over all experiments. Shading indicates ± SEM. **(E)** Cumulative response time profiles of AWA responses to AWA::Chrimson stimulation in WT versus *unc-13* animals. **(F)** Cumulative response time profiles of AWA and AIA responses to AWA::Chrimson stimulation in *unc-13* animals. Arrow indicates the delay between the time at which 50% of AWA versus 50% of AIA neurons have responded. **(G)** Delay between the time at which 50% of AWA versus 50% of AIA neurons responded to 1.15 µM diacetyl and AWA::Chrimson stimulation in WT and *unc-13* animals. WT responses are the same as in Figure 1. Bars are mean ± SEM. Asterisks refer to Kolmogorov-Smirnov test significance versus WT over full 10 s stimulus pulse. ns: not significant; *: p<0.05; ***: p<0.001. See Table S2 for sample sizes and test details.

The increased reliability of AIA responses in *unc-13* and *unc-18* mutants was even more striking when AWA was stimulated with AWA::Chrimson (Figures 3B and 3C, Figure 3 – figure supplement 1B-C). In the synaptic mutants, half of the AIA neurons responded to AWA::Chrimson within 1.2 seconds, a dramatic decrease in latency compared to wild type (>6 seconds) (Figure 3C). The AWA-to-AIA delay in synaptic mutants after AWA::Chrimson stimulation matched the delay observed to high concentrations of diacetyl (Figures 3F and 3G).

The magnitude and dynamics of individual AIA responses were unchanged in synaptic mutants compared to the wild type, regardless of whether they were elicited by diacetyl or AWA::Chrimson (Figure 3D, Figure 3 – figure supplement 1D-G). Control experiments demonstrated that direct AWA responses to optogenetic stimulation did not increase in *unc-13* synaptic mutants (Figure 3E, Figure 3 – figure supplement 1H-I). Chemical synapses thus inhibit AIA interneurons primarily by decreasing the reliability of AIA’s response at a given level of AWA activation.

### Glutamatergic sensory neurons cooperate to inhibit AIA

Eighteen pairs of neurons form chemical synapses onto AIA, six of which are sensory neurons that use glutamate as a neurotransmitter (White et al., 1986; Cook et al., 2019; Serrano-Saiz et al., 2013). Glutamate hyperpolarizes AIA, likely by activating glutamate-gated chloride channels, so these glutamatergic sensory neurons are plausible sources of the synaptic inhibition of AIA (Chalasani et al., 2010; Shinkai et al., 2011; López-Cruz et al., 2019). We selectively inhibited glutamate release using a CRISPR-edited version of the endogenous *eat-4* locus, which encodes the major vesicular glutamate transporter in *C. elegans*, in combination with cell-specific excision with flippase recombinase (Lee et al., 1999; López-Cruz et al., 2019) (Figures 4A and 4B). Selective excision of *eat-4* in four *tax-4-*expressing sensory neurons, AWC, ASE, ASK, and ASG, allowed AWA::Chrimson stimulation to evoke reliable AIA responses similar to those in *unc-18* synaptic transmission mutants (Figure 4C). This effect was not observed with either the modified *eat-4* locus or flippase expression alone (Figure 4 – figure supplement 1).

**Figure 4.**
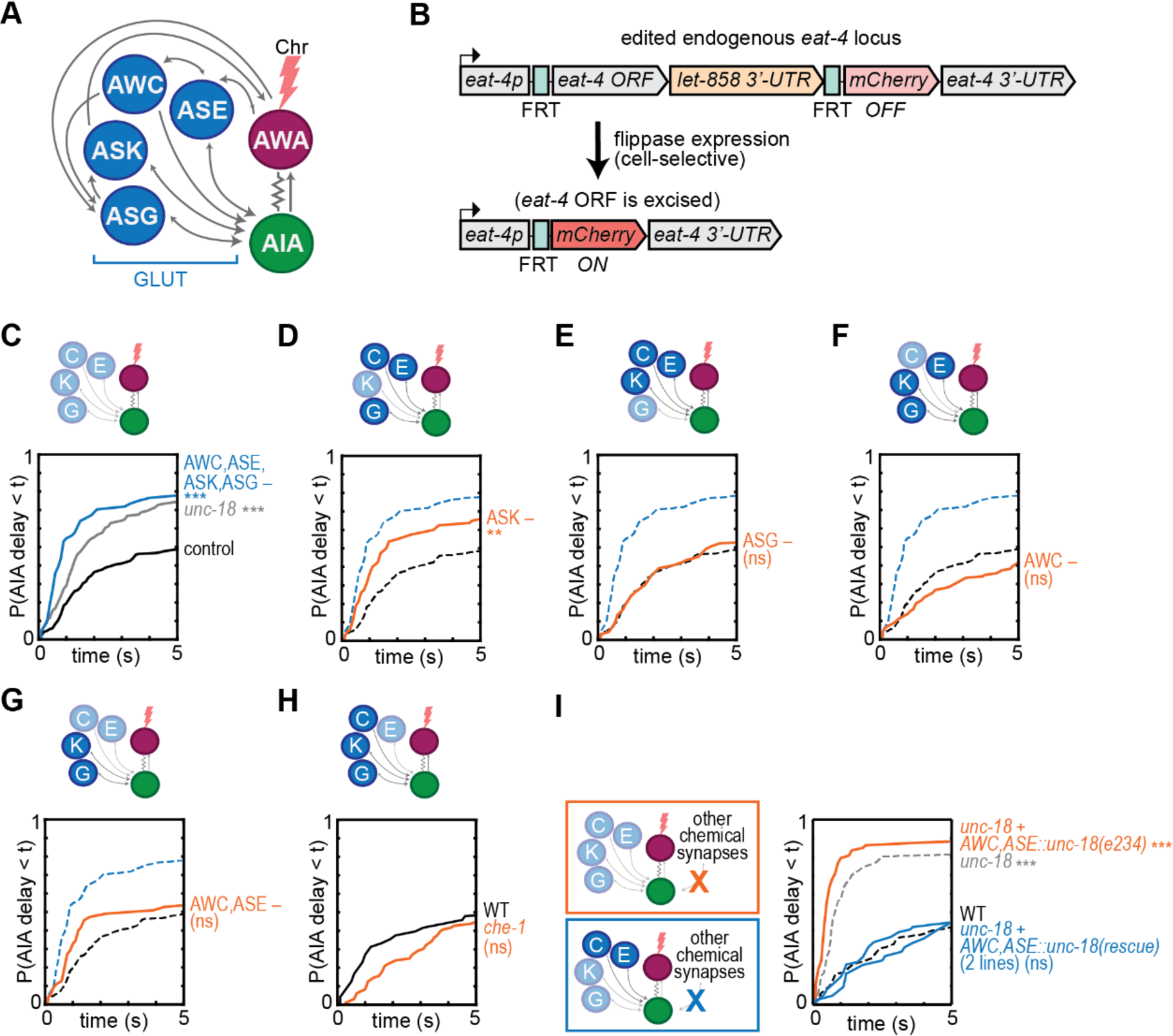
Glutamatergic sensory neurons cooperate to inhibit AIA. (A) Simplified diagram of connections between AWA, AIA and four glutamatergic sensory neurons, based on White et al. (1986). **(B)** Schematic of cell-selective glutamate knockout genetic strategy (López-Cruz et al, 2019). The *eat-4* locus is excised only in the presence of flippase. ORF: open reading frame; UTR: untranslated region; FRT: flippase recombinase target.**(C-H)** Cumulative response time profiles of AIA responses to AWA::Chrimson stimulation in various animals lacking either glutamate release or cellular function of specific sensory neurons. For (D-G), dotted black and blue lines are control and *eat-4-FRT; AWC+ASE+ASK+ASG::nFlippase*, respectively, from (C). **(C)** Control (*eat-4-FRT* genetic background with no flippase expression), *unc-18*, and *eat-4-FRT; AWC+ASE+ASK+ASG::nFlippase* animals. **(D)** *eat-4-FRT; ASK::nFlippase* animals. **(E)** *eat-4-FRT; ASG::nFlippase* animals. **(F)** *eat-4-FRT; AWC::nFlippase* animals. **(G)** *eat-4-FRT; AWC+ASE::nFlippase* animals. **(H)** WT and *che-1* animals (ASE cell fate mutants). (I) Cumulative response time profiles of AIA responses to AWA::Chrimson stimulation in WT, *unc-18* animals, *unc-18; AWC+ASE::unc-18(+)* transgenic animals (two lines), and *unc-18; ASE+AWC*::*unc-18(e234)* control transgenic animals. Asterisks refer to Kolmogorov-Smirnov significance versus *eat-4-FRT* controls (C-G) or WT (H, I) over full 10 s stimulus pulse. ns: not significant; **: p<0.01; ***: p<0.001. See Table S2 for sample sizes and test details.

**Figure 5.**
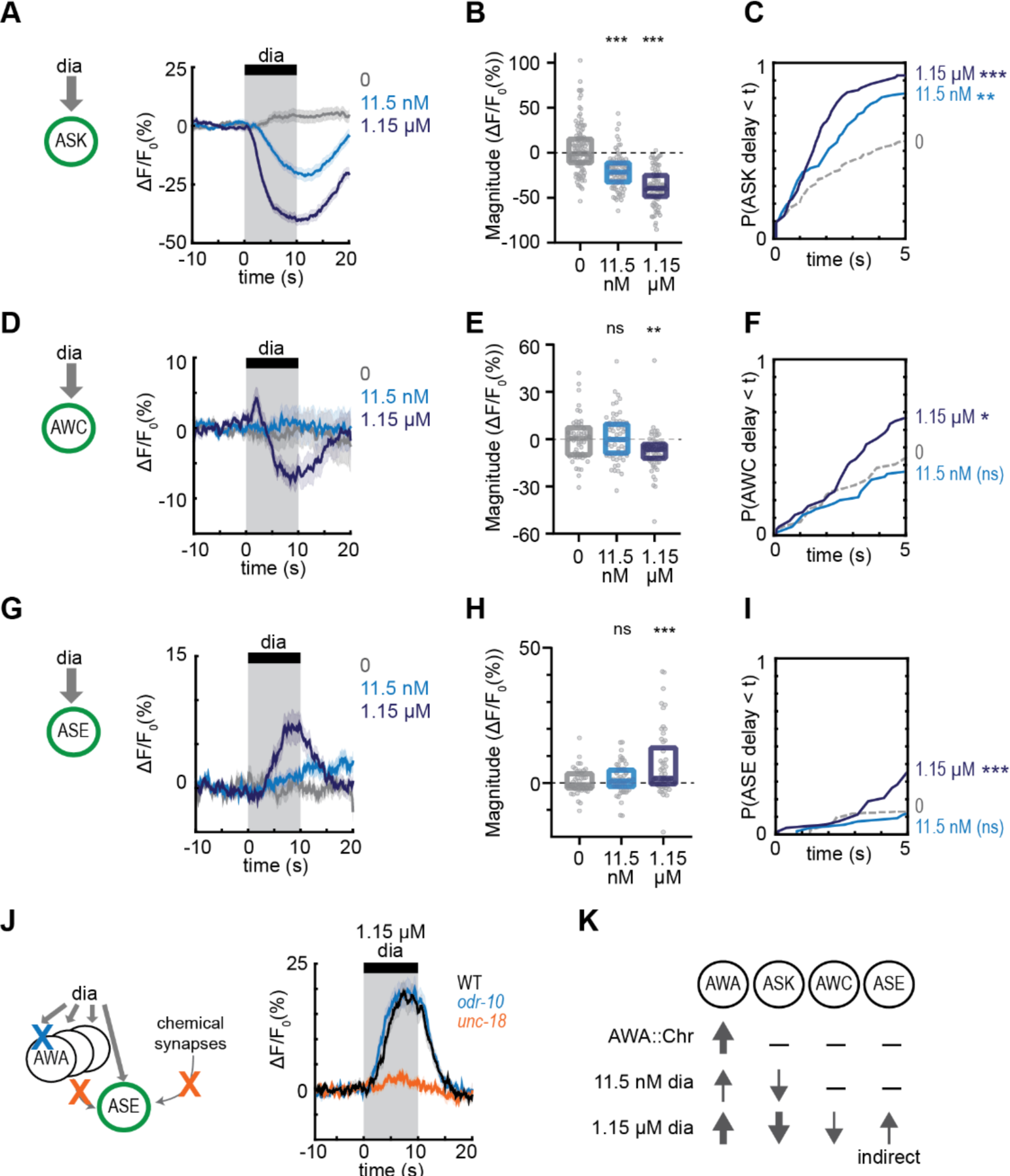
Multiple sensory neurons detect diacetyl. **(A, D, and G)** Mean ASK (A), AWC (D), and ASE (G) responses to 10 s pulses of buffer (0) or 11.5 nM or 1.15 µM diacetyl. Shading indicates ± SEM. **(B, E, and H)** Magnitude of individual responses from (A), (D), and (G). Boxes show median and interquartile range. **(C, F, and I)** Cumulative response time profiles of responses from (A), (D), and (G). Only the first 5 seconds are shown. **(J)** Mean ASE responses to 1.15 µM diacetyl in WT versus *unc-18* animals (synaptic transmission mutants) and *odr-10* animals (AWA diacetyl receptor mutants). ASE responses in *unc-18* animals are greatly diminished. Shading indicates ± SEM. **(K)** Summary of sensory neuron responses to various stimuli. Upward arrows indicate activation; downward arrows indicate inhibition. Arrow thickness reflects response magnitude. For (B), (E), and (H), asterisks indicate statistical significance of one-way ANOVA with Dunnett’s multiple comparisons test. ns: not significant; **: p<0.01; ***: p<0.001. See Table S3 for sample sizes and test details. For (C), (F), and (I), asterisks refer to Kolmogorov-Smirnov test significance versus buffer over full 10 s stimulus pulse. ns: not significant; *: p<0.05; **: p<0.01; ***: p<0.001. See Table S2 for sample sizes and test details.

No individual sensory pair accounted for the full effect of preventing glutamate release from AWC, ASE, ASK, and ASG together (Figures 4D-4H), although a significant partial effect was observed upon inhibition of glutamate release from ASK alone (Figure 4D). As previous studies implicated AWC and ASE in diacetyl responses (Larsch et al., 2015), we further examined the effect of chemical synapses from these two neurons. As noted above, the *unc-18* mutant has increased reliability of AIA responses to AWA::Chrimson stimulation, but in two independent lines in which *unc-18* was selectively rescued in ASE and AWC, AIA interneuron responses were restored to unreliable responses resembling wild type (Figure 4I). This effect was not observed with a control transgene that encoded the inactive *unc-18(e234)* mutant form in AWC and ASE. Synaptic vesicle release from AWC and ASE is therefore sufficient to inhibit AIA activation by AWA. In summary, multiple sensory neurons, including ASK and at least one of AWC and ASE, can release glutamate to inhibit AIA activation.

### Multiple sensory neurons detect diacetyl

To explain these results, we hypothesized that glutamatergic sensory neurons tonically inhibit AIA, and are inhibited when diacetyl is present to disinhibit AIA. This hypothesis agrees with previous calcium imaging studies showing that ASK and AWC are active at rest and inhibited by amino acids, pheromones, and certain odors (Wakabayashi et al., 2009; Macosko et al., 2009; Zaslaver et al., 2015; Chalasani et al., 2007; Kato et al., 2014). To extend this observation to diacetyl, we expressed the genetically-encoded calcium indicator GCaMP5A in ASK and AWC, and GCaMP3 in ASE, and determined that ASK and AWC were inhibited by the addition of 1.15 µM diacetyl, whereas ASE was activated by diacetyl at the same concentration (Figures 5A-5I). All responses were dose-dependent, with responses that were weaker (ASK) or absent (AWC, ASE) at 11.5 nM diacetyl (Figures 5A-5I). ASH, another glutamatergic sensory neuron that forms chemical synapses onto AIA, did not respond to 1.15 µM diacetyl (Figure 5 – figure supplement 1A-B), demonstrating neuronal specificity of the response.

*C. elegans* sensory neurons are abundantly interconnected by chemical synapses (White et al., 1986). Diacetyl responses in ASK and AWC were unchanged in *unc-18* synaptic transmission mutants, suggesting that these neurons detect diacetyl directly (Figure 5 – figure supplement 1D-G). By contrast, diacetyl responses in ASE were eliminated in *unc-18* mutants, suggesting that they are indirectly relayed from another sensory neuron (Figure 5J, Figure 5 – figure supplement 1C). AWA was not the source of the diacetyl response in ASE, as the response was preserved in an *odr-10* diacetyl receptor mutant (Figure 5J, Figure 5 – figure supplement 1C). Neither ASK, AWC, nor ASE showed a calcium response upon AWA::Chrimson stimulation (Figure 5 – figure supplement 1H-I).

In summary, AIA responses are unreliable upon AWA::Chrimson stimulation, which activates only AWA; more reliable to 11.5 nM diacetyl, which activates AWA and inhibits ASK; and highly reliable to 1.15 µM diacetyl, which activates AWA and ASE and inhibits AWC and ASK, leading to disinhibition of AIA.

### Combinatorial activation of AIA by isoamyl alcohol-sensing neurons

Multiple sensory neurons responded to diacetyl and together determined the reliability of the AIA response. We wondered whether the same logic applied for other odors. Based on previous ablation studies, chemotaxis to diacetyl requires AWA at low concentrations, with a redundant role for AWC at high concentrations (Chou et al., 2001), whereas chemotaxis to another odor and bacterial metabolite, isoamyl alcohol, requires AWC with a minor contribution from AWA (Bargmann et al., 1993; Worthy et al., 2018). Both AWC and AWA respond to isoamyl alcohol with calcium transients (Larsch et al., 2013), suggesting that study of this second odor could test the generality of the AIA activation model.

Previous work showed that AWC is inhibited by 9 and 90 µM isoamyl alcohol (Larsch et al., 2013; Yoshida et al., 2012; Gordus et al., 2015). We found that AWC was also inhibited by 0.9 µM isoamyl alcohol, and that AWA was activated by isoamyl alcohol in a graded fashion at 0.9, 9, and 90 µM isoamyl alcohol (Figures 6A, 6B, Figure 6 – figure supplement 1G-K). ASK was not inhibited by isoamyl alcohol at any tested concentration (Figure 6C, Figure 6 – figure supplement 1C, F).

**Figure 6.**
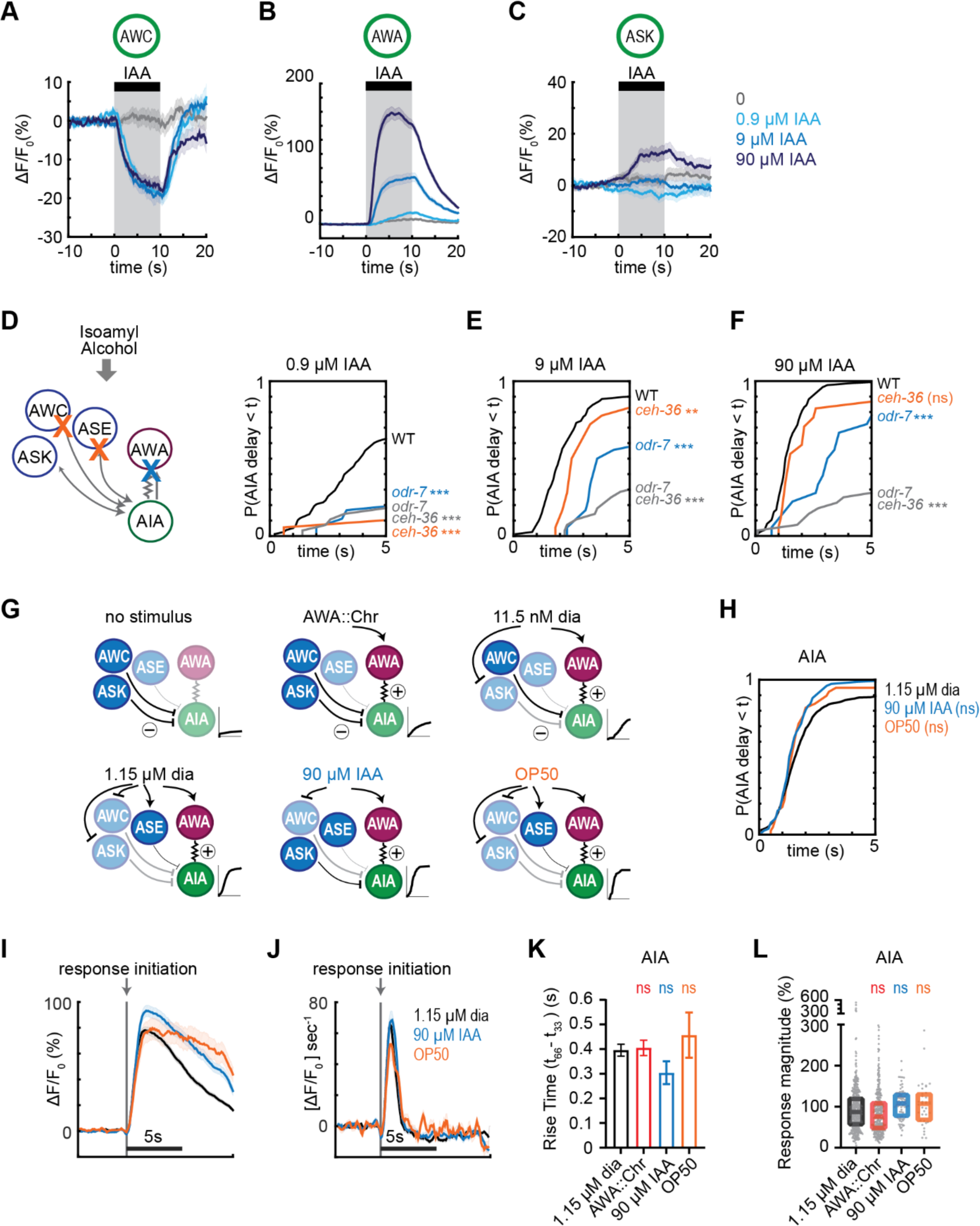
Combinatorial activation of AIA by isoamyl alcohol-sensing neurons. **(A – C)** Mean AWC (A), AWA (B), and ASK (C) sensory responses to 0, 0.9, 9, and 90 µM isoamyl alcohol in WT animals. Shading indicates ± SEM. **(D – F)** Cumulative response time profiles of AIA responses to 0.9 µM (D), 9 µM (E), and 90 µM (F) isoamyl alcohol in WT versus *odr-7* animals (AWA cell fate mutants), *ceh-36* animals (AWC and ASE cell fate mutants), and *odr-7 ceh-36* animals. **(G)** Summary of sensory neuron and AIA responses to different stimuli. Arrows represent activation, blunt lines represent inhibition. **(H)** Cumulative response time profiles of AIA responses to 1.15 µM diacetyl, 90 µM isoamyl alcohol, and *E. coli* OP50 bacteria-conditioned medium. **(I and J)** Mean AIA responses (I) and associated time derivatives (J) from (H) that initiated within 5 s of stimulus, aligned to the frame at which activation began. Shading indicates ± SEM. 1.15 µM diacetyl: n = 390; 90 µM isoamyl alcohol: n = 76; OP50: n = 37. **(K)** Rise time of responses from (I), with AIA responses to AWA::Chrimson stimulation from Figure 1I shown for comparison. Bars indicate SEM. **(L)** Magnitude of responses from (H), omitting traces that did not produce a detectable response, with AIA responses to AWA::Chrimson stimulation from Figure 1H. Boxes show median and interquartile range. For (D-F), asterisks refer to Kolmogorov-Smirnov test significance versus WT over full 10 s stimulus pulse. ns: not significant; *: p<0.05; **: p<0.01; ***: p<0.001. See Table S2 for sample sizes and test details. For (H), ns signifies a lack of significance of Kolmogorov-Smirnov test versus 1.15 µM diacetyl. See Table S2 for sample sizes and test details. For (K) and (L), ns signifies a lack of significance of one-way ANOVA with Dunnett’s multiple comparisons test versus 1.15 µM diacetyl. See Table S3 for sample sizes and test details for (L); see Table S4 for sample sizes and test details for (K).

We then examined AIA interneuron responses to 0.9, 9, and 90 µM isoamyl alcohol. As with diacetyl, AIA calcium responses were unreliable at the lowest concentrations, but became more reliable as isoamyl alcohol concentration increased (Figures 6D-6F). The lag between AWC and AIA responses also decreased as isoamyl alcohol concentration increased (Figure 6 – figure supplement 1K). At all concentrations, AWC responded first and AIA responded later, near the time of AWA activation (Figure 6 – figure supplement 1H-K). Interestingly, the opposite pattern held for diacetyl, where AWA responded first and AIA responded later, near the time of ASK inhibition (Figure 6 – figure supplement 1G).

To test the contributions of different sensory neurons to AIA activation by isoamyl alcohol, we monitored AIA responses in wild type, AWA cell fate mutants (*odr-7*), AWC and ASE cell fate mutants (*ceh-36*) (Lanjuin et al., 2003), and *odr-7 ceh-36* double mutants. *odr-7* mutants had unreliable AIA responses to isoamyl alcohol at all concentrations (Figures 6D-6F). At the higher concentrations of 9 µM and 90 µM isoamyl alcohol, AIA responses were more reliable in the *odr-7* and *ceh-36* single mutants than in the *odr-7 ceh-36* double mutants, indicating that AWA and AWC are partly redundant for AIA activation. At the lower concentration of 0.9 µM isoamyl alcohol, only the wild type animals had reliable AIA responses, indicating that both AWA and AWC are independently required for AIA activation.

We also examined the AIA response to a more complex food stimulus, *Escherichia coli* OP50-conditioned medium. This stimulus activates AWA and ASE and inhibits AWC, ASK, and several other sensory neurons (Zaslaver et al., 2015). *E. coli*-conditioned medium elicited reliable AIA calcium responses with similar latency, magnitude, and dynamics to those elicited by diacetyl and isoamyl alcohol (Figures 6G-6L).

## Discussion

### AIA uses AND-gate logic to integrate sensory information

Our work defines the contributions of individual sensory neurons to the activation of a downstream integrating neuron. Multiple *C. elegans* sensory neurons respond to an attractive odor such as diacetyl or isoamyl alcohol with cell type-specific dynamics and signs. The sensory neurons signal the presence of odor to AIA using electrical synapses (AWA) or glutamatergic chemical synapses (AWC, ASK). AIA does not respond reliably to input from a single sensory neuron. Rather, AIA is activated by coordinated inputs from multiple sensory neurons, functioning as an AND-gate that requires both activation by AWA, and disinhibition from glutamatergic neurons including AWC and ASK.

The AND-gate is an established motif in transcriptional regulation (Buchler et al., 2003), and bears similarity to the related concept of coincidence detection in neural circuits (Koch, 1999). In contrast with computational models of neurons based on additive inputs onto the target neuron (Dayan and Abbott, 2001; McCulloch and Pitts, 1943), the AND-gate computation is nonlinear and multiplicative (Koch, 1999). In the context of AIA, this logical computation requires multiple sensory neurons to report the presence of an attractant, an integrative step that may filter out environmental noise. Based on our results with diacetyl and isoamyl alcohol, it appears that different combinations of sensory neurons can generate the disinhibition and excitation necessary for reliable AIA activation.

It is interesting to compare AIA to another well-characterized interneuron, AIB. AIB activity rises when odors such as isoamyl alcohol are removed, triggering aversive behaviors such as reversals and turns (Chalasani et al., 2007). The coupling of odors to AIB activity is less reliable than coupling to AIA, and the mechanism of activation differs at a circuit level: AIB integrates sensory input with motor commands, such that its response to odor depends on the downstream motor state (Gordus et al., 2015). The wiring diagram predicts that AIA and AIB receive synaptic input from many of the same sensory neurons, but while AIA receives little feedback from interneurons, AIB has gap junctions and chemical synapses with multiple interneurons that drive motor states (White et al., 1986). These wiring patterns may explain why AIA activity is more closely coupled to sensory state, and less to motor state, than AIB. That said, AIB sensory integration has not been examined at the single neuron resolution used here, and integrated sensory state might also contribute to AIB activation.

### Excitation and disinhibition by electrical and chemical synapses

In *C. elegans,* behavioral functions of many neurons are known – for example, AIA supports forward movement in the presence of odor, reversals upon odor removal, and aversive olfactory learning – but the associated synaptic computations are not as well understood. As predicted by the wiring diagram, our results indicate that AWA primarily excites AIA via an electrical synapse, in the context of chemical synapses from other neurons. Mixed chemical-electrical synapses are present in many escape circuits, including insect giant fibers, goldfish Mauthner cells, and the *C. elegans* backward command neurons (Allen and Murphey, 2007, Phelan et al., 2008, Liu et al., 2017). In these escape circuits, the same cells form chemical and electrical synapses with each other to maximize speed and efficiency. By contrast, AIA receives chemical and electrical synapses from different sensory neurons to enable integration.

The AWA-AIA gap junctions preferentially transmit excitatory signals from AWA to AIA. In *C. elegans*, as in *Drosophila* and vertebrates, asymmetrical electrical synapses can form at heterotypic or heteromeric gap junctions (Liu et al., 2013; Miller et al., 2017; Phelan et al., 2008). Consistently, AWA and AIA express distinct sets of innexin genes (Bhattacharya et al., 2019). The chemical synapses onto AIA are inhibitory and glutamatergic. Direct electrophysiological measurements indicate that glutamate inhibits AIA, likely via glutamate-gated chloride channels (López-Cruz et al., 2019). Behavioral studies have also demonstrated glutamatergic inhibition of AIA, both through the glutamate-gated chloride channel GLC-3 and through the metabotropic glutamate receptor MGL-1 (Shinkai et al., 2011; Chalasani et al., 2010; López-Cruz et al., 2019).

At a synaptic level, the combination of excitatory and disinhibitory inputs of AIA resemble interactions at the glomeruli of the rat inferior olivary nucleus (Pereda et al., 2013). Principal cells in the inferior olive are coupled via electrical synapses, and are uncoupled when neurons from the deep cerebellar nuclei release GABA at adjacent chemical synapses. The inhibitory synapses shunt excitatory current flow at the electrical synapse (Pereda et al., 2013). Synchronized firing in this system has features of an AND-gate, with distinct states that depend on both electrical and chemical synapses.

At a circuit level, the roles of excitation and inhibition in an AND-gate differ from their roles in many circuits. Excitation and inhibition are most frequently observed as balanced inputs, tightly correlated in time and space, poising circuits to detect small changes with precision (Okun and Lampl, 2008; Denève and Machens, 2016). In the AND-gate logic employed by AIA, excitation and inhibition alternate, switching the neuron’s activity state. A related disinhibition logic, with a more complex mechanism, is used in mammalian cortical and hippocampal circuits, where VIP-expressing interneurons disinhibit information flow through excitatory circuits by GABAergic inhibition of local inhibitory interneurons (Pi et al., 2013; Turi et al., 2019).

### AIA activity as a readout of positive valence

We suggest that AIA signals that a given stimulus is worth pursuing, based on an integrated sensory state that reflects the positive valence and stability of the stimulus. The integrated positive valence is represented by the activity state of multiple sensory neurons. The relevant combinations of sensory neurons vary, and are partly distinct from the roles of those sensory neurons in chemotaxis. For example, AWA is not necessary for chemotaxis to isoamyl alcohol, but has an important role in AIA activation by that odor, and the converse holds for ASK and diacetyl. Further evidence for combinatorial integration by AIA comes from behavioral studies in which animals cross an aversive copper barrier, sensed by the glutamatergic ASH sensory neurons, to reach an attractive diacetyl source (Shinkai et al., 2011). The copper-diacetyl behavioral interaction requires ASH glutamatergic inhibition of AIA via the glutamate-gated chloride channel GLC-3. In this context, the integrated sensory state detected by AIA includes a repellent as well as an attractant, and a different combination of sensory inputs.

A second feature of AIA activation is a delay that suggests that the AND-gate filters out transient or noisy odor stimuli. For both diacetyl and isoamyl alcohol, AIA activation coincides with the slower of the excitatory or disinhibitory sensory neurons. Because sensory neurons have different dynamics, in most cases AIA activation would require the sustained presence of odor. A requirement for sustained stimulus can also reside in the sensory neurons themselves, as exemplified by the one-second delay between stimulus onset and an AWA action potential (Liu et al., 2018). This requirement for a sustained input contrasts with the fast sampling that optimizes steering decisions during chemotaxis (Iino and Yoshida, 2009; Kato et al, 2014). For both diacetyl and isoamyl alcohol, the first neuron to respond to the odor (AWA for diacetyl, AWC for isoamyl alcohol) is the neuron that is most important for chemotaxis behavior.

At a single trial level, AIA calcium dynamics and magnitudes are stereotyped, whether activated by individual odors, optogenetic stimulus, or by *E. coli* conditioned medium; only the probability and the latency vary. The stereotyped responses suggest that AIA is not a passive transmitter of sensory information. Instead, it compresses varied sensory dynamics into a uniform and time-limited signal of a salient positive stimulus. Its integrative quality and slower temporal features may allow AIA to provide a working memory of positive chemosensory environments.

## Materials and Methods

### *C. elegans* Growth

We used standard genetic and molecular techniques (Brenner, 1974). *C. elegans* strains were maintained at 22°C on Nematode Growth Media (NGM; 51.3 mM NaCl, 1.7% agar, 0.25% peptone, 1 mM CaCl_2_, 12.9 µM cholesterol, 1 mM MgSO_4_, 25 mM KPO_4_ at pH 6) plates seeded with LB-grown *Escherichia coli* OP50 bacteria. Animals had constant access to food for at least 3 generations prior to experiments. Experiments were performed on young adult hermaphrodites. All strains are listed in Table S5.

### Stimulus Preparation

Stimulus solutions were freshly prepared each experimental day by serially diluting from a pure stock of diacetyl (2,3-butanedione; Sigma-Aldrich 11038, CAS 431-03-8; stored at 4°C) or isoamyl alcohol (EMD AX-1440-6, CAS 123-51-3; stored at 4°C), or by directly dissolving NaCl (Fisher Chemical S271-1, CAS 7647-14-5) into S Basal buffer (0.1 M NaCl, 5.74 mM K_2_HPO_4_ and 44.1 mM KH_2_PO_4_ at pH 6, 5 µg/ml cholesterol). *E. coli* OP50 conditioned medium was prepared by seeding 30 ml NGM buffer without agar or cholesterol with a colony of OP50 bacteria and shaking the culture at 37°C overnight such that optical density was ∼2 by the morning of the experiment. The OP50 culture was filtered (0.22 µm Millex GP) before the experiment and NGM was used as the buffer control for these experiments. All stimulus solutions contained 1 mM (-)-tetramisole hydrochloride (Sigma L9756, CAS 16595-80-5) to paralyze the body wall muscles and were stored in brown glass vials.

### Calcium Imaging

The calcium imaging protocol was adapted from Larsch et al. (2015). Larvae expressing GCaMP were selected as L4s the evening before the experiment and picked to new OP50 plates. For Chrimson experiments, L4s were transferred animals to freshly seeded plates of 5x concentrated OP50-seeded LB with or without 5 µM all-trans retinal (Sigma R2500; CAS 116-31-4) and housed animals overnight in complete darkness.

Before beginning the experiment, we selected animals for visible GCaMP fluorescence and gently washed them in S Basal buffer. We then loaded ∼10 animals of two genotypes or conditions into separate arenas of a custom-fabricated two-arena polydimethylsiloxane (PDMS; Sigma 761036, made from 9:1 base:curing agent, Sylgard 184) imaging devices that had been de-gassed in a vacuum dessicator for at least 5 minutes. We chose ∼10 animals per arena to maximize the number of replicates within a condition or genotype without sacrificing proper flow. Animals were paralyzed in darkness for ∼90 minutes in buffer + tetramisole before the start of the recording.

Experiments were performed on a Zeiss AxioObserver A1 inverted microscope fit with a 5x/0.25 NA Zeiss Fluar objective. A Hammamatsu Orca Flash 4 sCMOS camera was mounted to the microscope using a 0.63x c-mount adapter to increase field of view. We delivered 474 nm wavelength light with a Lumencor SOLA-LE lamp at 165 mW/cm^2^ for odor-only experiments and at 40 mW/cm^2^ for experiments involving optogenetics to avoid blue light activation of the Chrimson channel. Videos were acquired at 10 fps (100 ms exposure time), with excitation light pulsed at a 10 ms per exposure duty cycle. We used Metamorph 7.7.6 software to control image acquisition and light pulsing in addition to rapid stimulus switching (National Instruments NI-DAQmx connected to an Automate Valvebank 8 II actuator that controls a solenoid valve), odor selection (Hamilton 8-way distribution valve), and activation of an external red LED for Chrimson stimulation (Mightex Precision LED Spot Light, 617 nm, PLS-0617-030-S; attached to Chroma ET605/50x filter to narrow band to 605 ± 25 nm). For Chrimson experiments, red light intensity was 15 mW/cm^2^.

Animals received two pulses of the tested stimulus, and both pulses were pooled for analyses, with the exception of Figure 1 – figure supplement 1I-Q. In experiments with multiple odor concentrations, we delivered odors in order of increasing concentration. In Chrimson experiments with diacetyl controls (Figure 1 – figure supplement 1F), or diacetyl experiments with NaCl controls (Figure 5 – figure supplement 1A-B), the control was delivered last. To confirm proper flow, we delivered a pulse of fluorescein dye at the end of the experiment; assays in which flow was impeded were discarded.

Raw fluorescence values were measured using a custom ImageJ script from Larsch et al. (2013), which measures the average intensity of a 4×4 pixel square and subtracts the local background intensity. For all sensory neurons, the square captured the soma; for AIA, it captured the middle of the neural process. Animals that moved too much for the tracking script, and animals with no visible AIA soma, were discarded. Occasionally, the tracking script inserted NaN instead of a fluorescence intensity; pulses with NaN values were discarded. We did not mask our data, although the tracking script is semi-automated to reduce experimenter bias.

Each background-subtracted raw fluorescence trace was first normalized to generate ΔF/F_0_, where F_0_ was the median of the 10 seconds (100 frames) before the odor pulse onset. Traces were then smoothed by 5 frames such that each frame *t* represented the mean of *t*-2 to *t*+2.

In a preliminary experiment, we delivered 28 AWA::Chrimson pulses to wild type animals, and 24 pulses to *unc-13(e51)* mutant animals (not included in analyses because we used a slightly different protocol). This experiment captured a statistically significant difference in cumulative response time profiles between wild type and *unc-13* AIA responses to AWA::Chrimson stimulation (Kolmogorov-Smirnov test, p: 0.007, D test statistic: 0.470). Guided by this result, all experiments reported here included at least 17 light or odor pulses (and typically >30) per genotype or condition.

We delivered two light or odor pulses per animal, tested 5-10 animals per PDMS device arena, tested 2-16 arenas per genotype or condition, and tested 2-10 genotypes or conditions per experimental block. The number of arenas per genotype or condition depended on the number of genotypes or conditions included in an experimental block, such that each genotype or condition was tested on at least two days with fresh odorant solutions or retinal plates, tested in both the upper and lower arenas of a PDMS device (see Figure 1 – figure supplement 1A), and tested simultaneously with the control at least once. The n used for analysis refers to individual stimulus pulses. Statistical analyses include all genotypes or conditions in an experimental block, corrected for multiple comparisons that in some cases include genotypes or conditions we do not present. The genotypes or conditions included in an experimental block depended on the hypothesis we were testing. We included a wild type-to-*unc-18(e234)* comparison in several independent experimental blocks, and the cumulative response time profiles for the separate and combined experimental blocks are shown in Figure 3 – figure supplement 1C.

### Determining Response Latency Times

For AWA, AIA, ASE, and ASH, a calcium trace was deemed a “response” at the first frame *t* at which the mean smoothed ΔF/F_0_ of *t* to *t*+12 exceeded 4 standard deviations of the mean of the 10 s pre-odor ΔF/F_0_, and the mean time derivative of *t* to *t*+1 exceeded 1 standard deviation of the mean of the 10 s pre-odor ΔF/F_0_.

To determine response latency, but not for other calculations, each ASK and AWC calcium trace was scaled such that the minimum value was 0 and the maximum value was 1. The calcium trace was deemed a “response” at the first frame *t* at which the mean scaled ΔF/F_max_ of *t* to *t*+10 was below 2 standard deviations of the mean of the 10 s pre-odor ΔF/F_0_, and either the time derivative of *t* to *t*+10 was below 0.5 standard deviations, or the time derivative to *t* to *t*+5 was below 1.15 standard deviations, of the mean of the 10 s pre-odor ΔF/F_0_.

To compare the variability of response latencies, we compared the cumulative response time profiles. We used the Kolmogorov-Smirnov test to compare these distributions since this test would capture both the latencies and probability of response. Although the figures show only 5 seconds of stimulus, the Kolmogorov-Smirnov test compared distributions for 10 seconds of stimulus. Details of each test, including the D test statistic, can be found in Table S2.

### Comparing AWA-to-AIA and AWC-to-AIA Lag Times

To calculate the mean lag between AWA, or AWC, and AIA responses, we subtracted the frame at which 50% of AWA or AWC neurons had responded from the frame at which 50% of AIA neurons had responded to a given stimulus. We performed this calculation 1000 times from randomly bootstrap-sampled populations that had the same *n* as the true population, sampled with replacement. The standard deviation of the bootstrapped distribution was used as the standard error of the bootstrapped mean.

### Measuring Magnitudes

To calculate ASK, AWC, ASE, and ASH response magnitudes to a given stimulus, we subtracted the mean ΔF/F_0_ of 10 frames (1 second) prior to stimulus delivery from the mean ΔF/F_0_ of the final 10 frames within the odor pulse. Because AWA and AIA responses often adapt during the stimulus pulse, we defined AWA and AIA response magnitudes as the maximum ΔF/F_0_ within the 10 s stimulus pulse.

To compare AIA or AWA magnitudes between genotypes or conditions, we included only calcium traces that represent detectable responses as described in the Determining Latency Times section. To determine whether there was an appreciable ASK, AWC, or ASH response to a given stimulus, or AWA response to isoamyl alcohol, we included all calcium traces.

We statistically tested differences with either an ordinary one-way ANOVA with Dunnett’s multiple comparisons test for experiments with more than two conditions, or an unpaired t-test for experiments with only two conditions. For ASK, AWC, ASE, AWA, and ASH responses to diacetyl, isoamyl alcohol, or NaCl, we used responses to S Basal – S Basal buffer switches as the control. For ASK, AWC, and ASE responses to AWA::Chrimson stimulation in Figure 5 – figure supplement 1I, we used a paired t-test to compare response magnitudes to the change in ΔF/F_0_ within a similar time window prior to the light pulse. Details of each test can be found in Table S3.

### Calculating Rise Times

To calculate rise times of AIA responses, we included only calcium traces that represent detectable responses as described in the Determining Latency Times section. We calculated the rise time by subtracting the time at which AIA reached 33% of its peak magnitude from the time at which AIA reached 66% of its peak magnitude for each response. We compared rise times to various stimuli using an unpaired t-test for Figure 1I, and an ordinary one-way ANOVA with Dunnett’s multiple comparisons test (3 comparisons) for Figure 6K.

## ACKNOWLEDGEMENTS

We thank Philip Kidd, Sagi Levy, Qiang Liu, Aylesse Sordillo, Elias Scheer, Du Cheng, Audrey Harnagel, Alejandro López-Cruz, Likui Feng, James Lee, Javier Marquina-Solis, and Andrew Gordus for thoughtful discussions and comments on the manuscript. We also thank Andrew Leifer, Vanessa Ruta, and Shai Shaham for scientific discussions about the work presented here. We thank Mei Zhen for the *unc-7 unc-9* double mutant strain. This work was supported by the Howard Hughes Medical Institute, of which CIB was an investigator, and by the Chan Zuckerberg Initiative.

## AUTHOR CONTRIBUTIONS

MD performed all experiments. MD and CIB designed experiments, interpreted data, and wrote the manuscript.

## COMPETING INTERESTS

The authors declare no competing financial interests.

## SUPPLEMENTAL TABLES

**Table S1.**
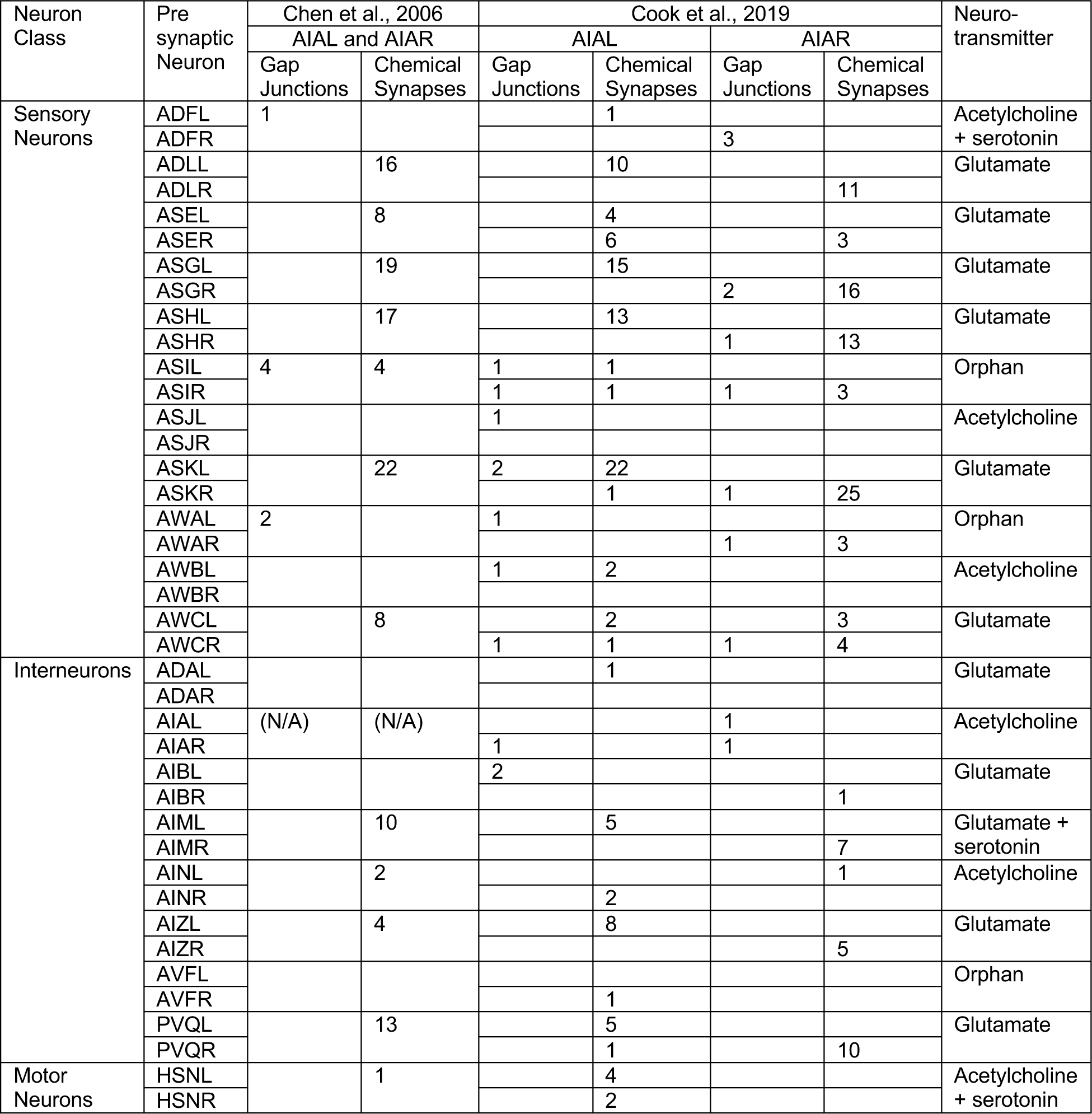
Presynaptic partners of AIA. Number of known chemical synapses onto AIA and gap junctions with AIA. Both Chen et al. (2006) and Cook et al. (2019) are based on same collection of serial-section electron micrographs from White et al. (1986). Neurotransmitter information is based on Pereira et al. (2015).

**Table S2.** Details of Cumulative Response Time Profiles. Kolmogorov-Smirnov test statistics and sample sizes for all cumulative response time profiles presented, calculated for full 10 s stimulus pulse. Italics indicate non-WT genetic backgrounds. D test represents the maximum effect size across the distributions. p-values below 0.05 are bolded for emphasis.

| Genotype or condition (test) | Genotype or condition (control) | Neuron Recorded | Stimulus | n (test) | n (control) | D test statistic | p-value | Figure |
| --- | --- | --- | --- | --- | --- | --- | --- | --- |
| AWA::Chr response to odor after light pulses | WT, no AWA::Chr expression or pulses | AIA | 1.15 μM dia | 90 | 438 | 0.174 | **0.022** | S1F |
| WT, 2<sup>nd</sup> pulse | WT, 1<sup>st</sup> pulse | AIA | AWA::Chr | 282 | 282 | 0.149 | **0.004** | S1I |
| WT, 2<sup>nd</sup> pulse | WT, 1<sup>st</sup> pulse | AIA | 11.5 nM dia | 115 | 115 | 0.105 | 0.709 | S1L |
| WT, 2<sup>nd</sup> pulse | WT, 1<sup>st</sup> pulse | AIA | 1.15 μM dia | 217 | 217 | 0.204 | **<0.001** | S1O |
| *odr-7(ky4)* | WT | AIA | 1.15 μM dia | 70 | 78 | 0.465 | **<0.001** | 2A |
| *odr-10(ky32)* | WT | AIA | 1.15 μM dia | 32 | 78 | 0.700 | **<0.001** | 2A |
| AWA::TeTx | WT | AIA | 1.15 μM dia | 27 | 20 | 0.243 | 0.508 | 2B |
| *unc-9(fc16) unc-7(e5)* | WT | AIA | 1.15 μM dia | 86 | 94 | 0.315 | **<0.001** | 2C |
| *unc-9(fc16) unc-7(e5)* | WT | AIA | AIA::Chr | 30 | 50 | 0.160 | 0.723 | 2E |
| *unc-9(fc16) unc-7(e5)* | WT | AWA | AIA::Chr | 48 | 58 | 0.096 | 0.968 | 2G |
| AWA::TeTx | WT | AIA | AWA::Chr | 56 | 146 | 0.158 | 0.264 | S2D |
| *unc-9(fc16) unc-7(e5)* | WT | AIA | 11.5 nM dia | 61 | 70 | 0.329 | **0.002** | S2G |
| *unc-9(fc16) unc-7(e5)* | WT | AIA | AWA::Chr | 96 | 146 | 0.204 | **0.016** | S2I |
| *unc-18(e234)* | WT | AIA | AIA::Chr | 46 | 64 | 0.108 | 0.914 | S2L |
| *unc-18(e234)* | WT | AWA | AIA::Chr | 69 | 58 | 0.343 | **0.001** | S2N |
| *unc-18(e234)* | WT | AIA | 11.5 nM dia | 34 | 66 | 0.642 | **<0.001** | 3A |
| *unc-13(e51)* | WT | AIA | 11.5 nM dia | 24 | 66 | 0.527 | **0.001** | 3A |
| *unc-18(e234)* | WT | AIA | AWA::Chr | 57 | 146 | 0.427 | **<0.001** | 3C |
| *unc-18(e81)* | WT | AIA | AWA::Chr | 40 | 146 | 0.415 | **<0.001** | 3C |
| *unc-13(e51)* | WT | AIA | AWA::Chr | 45 | 146 | 0.321 | **0.002** | 3C |
| *unc-13(e51)* | WT | AWA | AWA::Chr | 36 | 74 | 0.088 | 0.992 | 3E |
| *unc-18(e234)* | WT | AIA | 1.15 μM dia | 56 | 145 | 0.290 | **0.002** | S3A |
| *unc-18(e81)* | WT | AIA | 1.15 μM dia | 72 | 145 | 0.107 | 0.640 | S3A |
| *unc-13(e51)* | WT | AIA | 1.15 μM dia | 46 | 145 | 0.180 | 0.208 | S3A |
| *unc-31(e928)* | WT | AIA | AWA::Chr | 50 | 146 | 0.089 | 0.931 | S3J |
| *unc-18(e234)* | *eat-4-FRT* | AIA | AWA::Chr | 154 | 214 | 0.264 | **<0.001** | 4C |
| *eat-4-FRT*; AWC+ASE+ASK+ASG::nFlippase | *eat-4-FRT* | AIA | AWA::Chr | 64 | 214 | 0.360 | **<0.001** | 4C |
| *eat-4-FRT*; ASK::nFlippase | *eat-4-FRT* | AIA | AWA::Chr | 80 | 214 | 0.224 | **0.006** | 4D |
| *eat-4-FRT*; ASG::nFlippase | *eat-4-FRT* | AIA | AWA::Chr | 67 | 214 | 0.062 | 0.989 | 4E |
| *eat-4-FRT*; AWC::nFlippase | *eat-4-FRT* | AIA | AWA::Chr | 123 | 214 | 0.129 | 0.147 | 4F |
| *eat-4-FRT*; AWC+ASE::nFlippase | *eat-4-FRT* | AIA | AWA::Chr | 55 | 214 | 0.188 | 0.090 | 4G |
| *che-1(p674)* | WT | AIA | AWA::Chr | 96 | 48 | 0.188 | 0.211 | 4H |
| *unc-18(e234)* | WT | AIA | AWA::Chr | 70 | 80 | 0.521 | **<0.001** | 4I |
| *unc-18(e234)*; AWC+ASE::unc-18(e234) | WT | AIA | AWA::Chr | 72 | 80 | 0.629 | **<0.001** | 4I |
| *unc-18(e234)*; AWC+ASE::unc-18(WT)#1 | WT | AIA | AWA::Chr | 42 | 80 | 0.115 | 0.860 | 4I |
| *unc-18(e234)*; AWC+ASE::unc-18(WT)#2 | WT | AIA | AWA::Chr | 74 | 80 | 0.088 | 0.928 | 4I |
| *eat-4-FRT* | WT | AIA | AWA::Chr | 214 | 149 | 0.081 | 0.605 | S4 |
| AWC+ASE+ASK+ASG::nFlippase | WT | AIA | AWA::Chr | 37 | 149 | 0.179 | 0.658 | S4 |
| WT | WT | ASK | 0 vs. 11.5 nM dia | 82 | 115 | 0.301 | **0.004** | 5C |
| WT | WT | ASK | 0 vs. 1.15 μM dia | 84 | 115 | 0.404 | **<0.001** | 5C |
| WT | WT | AWC | 0 vs. 11.5 nM dia | 60 | 52 | 0.096 | 0.959 | 5F |
| WT | WT | AWC | 0 vs. 1.15 μM dia | 60 | 52 | 0.276 | **0.029** | 5F |
| WT | WT | ASE | 0 vs. 11.5 nM dia | 54 | 42 | 0.220 | 0.220 | 5I |
| WT | WT | ASE | 0 vs. 1.15 μM dia | 53 | 42 | 0.307 | **0.024** | 5I |
| *odr-7(ky4)* | WT | AIA | 0.9 μM IAA | 18 | 78 | 0.539 | **<0.001** | 6D |
| *ceh-36(ky640)* | WT | AIA | 0.9 μM IAA | 18 | 78 | 0.624 | **<0.001** | 6D |
| *odr-7(ky4) ceh-36(ky640)* | WT | AIA | 0.9 μM IAA | 28 | 78 | 0.571 | **<0.001** | 6D |
| *odr-7(ky4)* | WT | AIA | 9 μM IAA | 18 | 78 | 0.658 | **<0.001** | 6E |
| *ceh-36(ky640)* | WT | AIA | 9 μM IAA | 18 | 78 | 0.457 | **0.004** | 6E |
| *odr-7(ky4) ceh-36(ky640)* | WT | AIA | 9 μM IAA | 28 | 78 | 0.739 | **<0.001** | 6E |
| *odr-7(ky4)* | WT | AIA | 90 μM IAA | 18 | 77 | 0.687 | **<0.001** | 6F |
| *ceh-36(ky640)* | WT | AIA | 90 μM IAA | 17 | 77 | 0.237 | 0.416 | 6F |
| *odr-7(ky4) ceh-36(ky640)* | WT | AIA | 90 μM IAA | 28 | 77 | 0.796 | **<0.001** | 6F |
| WT | WT | AIA | 1.15 μM dia vs. 90 μM IAA | 78 (IAA) | 438 (dia) | 0.131 | 0.203 | 6H |
| WT | WT | AIA | 1.15 μM dia vs. OP50-conditioned medium | 39 (OP50) | 438 (dia) | 0.147 | 0.420 | 6H |
| WT | WT | AWC | 0 vs. 0.9 μM IAA | 42 | 42 | 0.429 | **<0.001** | S6D |
| WT | WT | AWC | 0 vs 9 μM IAA | 42 | 42 | 0.548 | **<0.001** | S6D |
| WT | WT | AWC | 0 vs 90 μM IAA | 42 | 42 | 0.691 | **<0.001** | S6D |
| WT | WT | AWA | 0 vs. 0.9 μM IAA | 78 (0) | 80 (0.9 μM) | 0.292 | **0.002** | S6E |
| WT | WT | AWA | 0 vs 9 μM IAA | 78 (0) | 80 (9 μM) | 0.558 | **<0.001** | S6E |
| WT | WT | AWA | 0 vs 90 μM IAA | 78 (0) | 80 (90 μM) | 0.772 | **<0.001** | S6E |
| WT | WT | ASK | 0 vs. 0.9 μM IAA | 60 | 60 | 0.510 | 0.510 | S6F |
| WT | WT | ASK | 0 vs 9 μM IAA | 60 | 60 | 0.133 | 0.660 | S6F |
| WT | WT | ASK | 0 vs 90 μM IAA | 60 | 60 | 0.167 | 0.375 | S6F |
| WT | WT | ASK vs. AIA | 1.15 μM dia | 84 (ASK) | 438 (AIA) | 0.128 | 0.202 | S6G |
| WT | WT | AWA vs. AIA | 0.9 μM IAA | 80 (AWA) | 438 (AIA) | 0.807 | 0.103 | S6H |
| WT | WT | AWA vs. AIA | 9 μM IAA | 80 (AWA) | 438 (AIA) | 0.677 | 0.115 | S6I |
| WT | WT | AWA vs. AIA | 90 μM IAA | 80 (AWA) | 438 (AIA) | 0.125 | 0.189 | S6J |

**Table S3.**
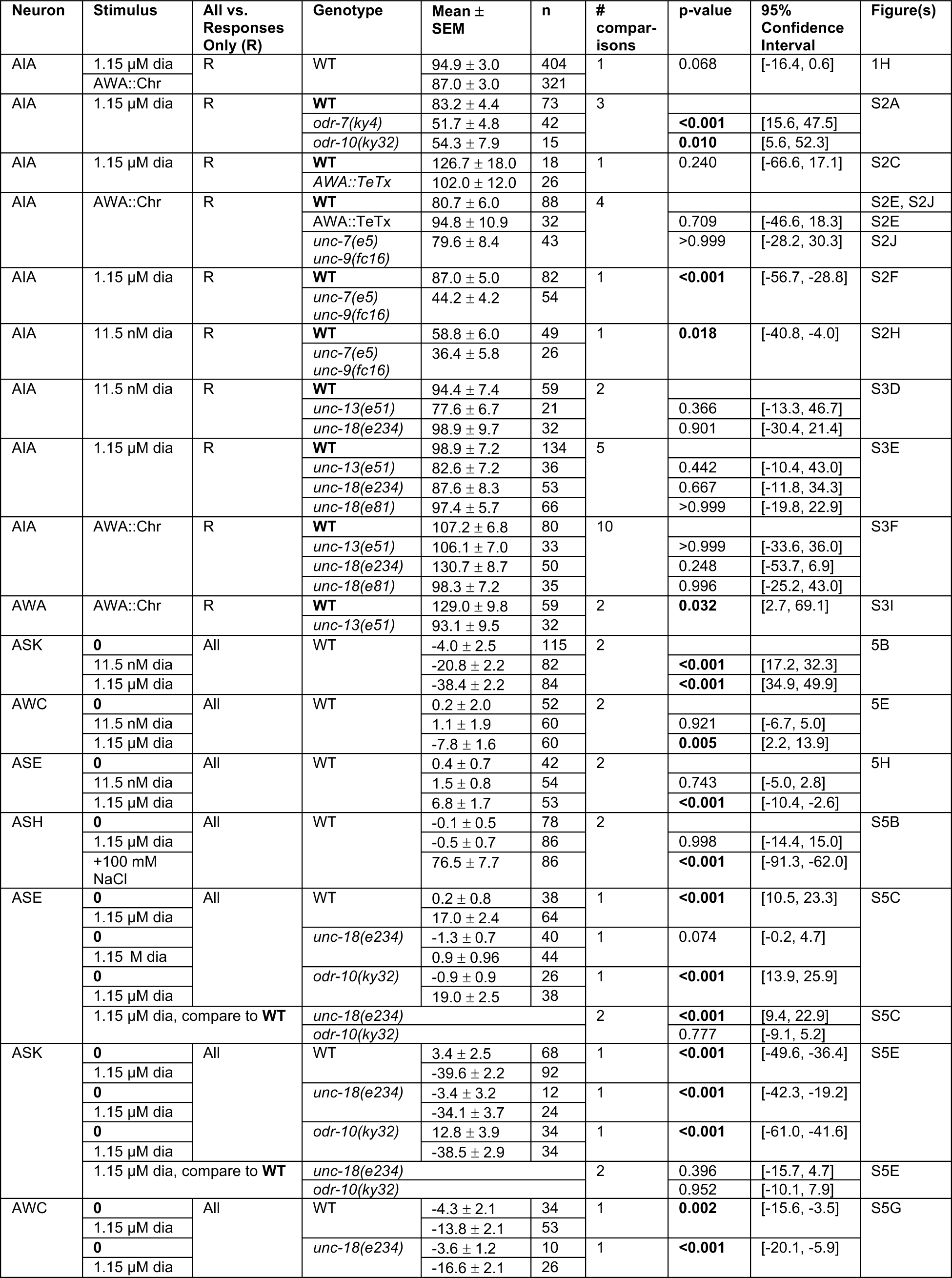
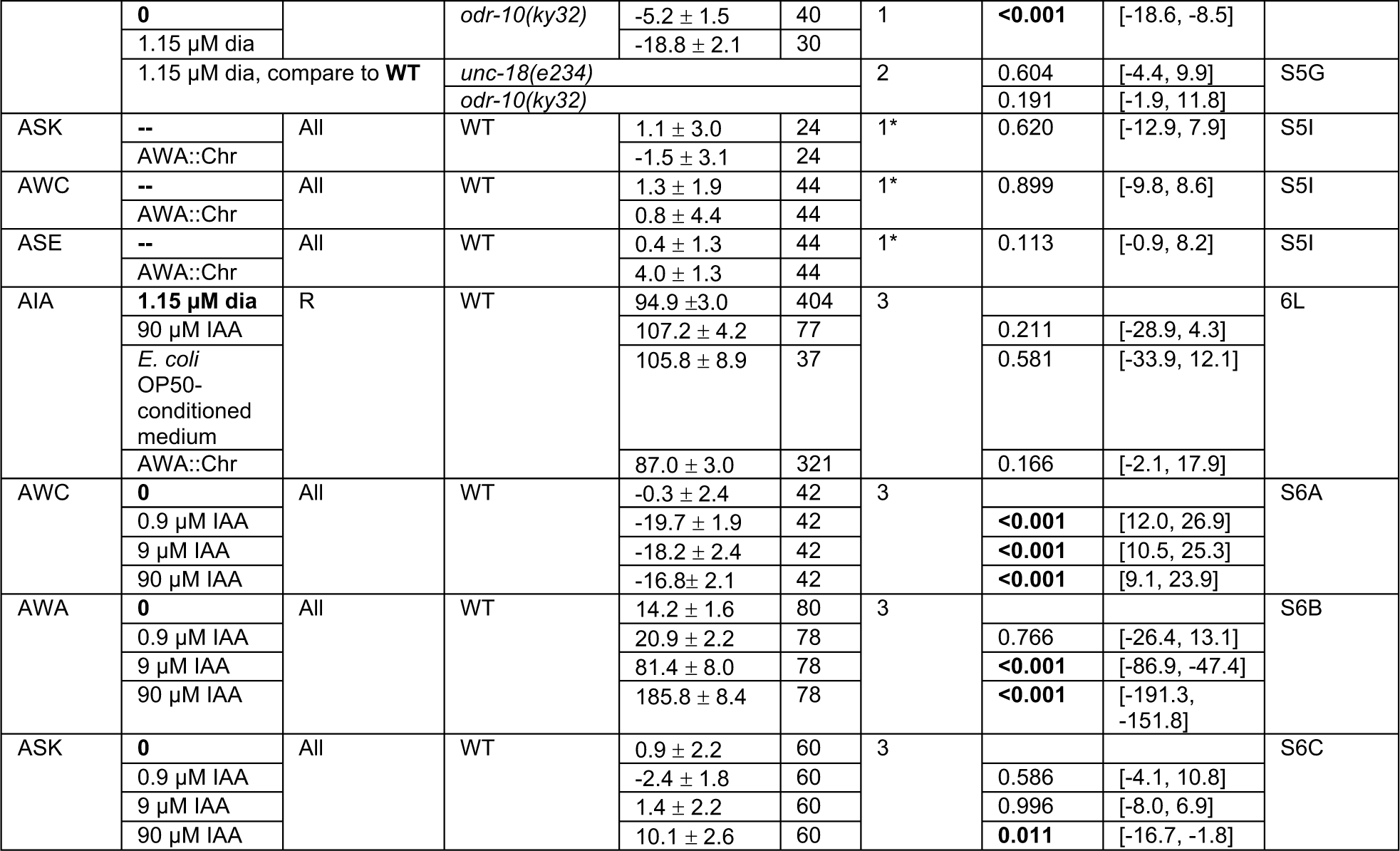
Details of Magnitude Comparisons. Magnitudes of responses to various stimuli, with either an unpaired t-test (if the number of comparisons is one) or an ordinary one-way ANOVA with Dunnett’s multiple comparisons test (if the number of comparisons exceeds one); * indicates paired t-test. Bolded genotype or stimulus indicates the control group used for comparisons. Italics indicate non-wildtype genetic background. p-values below 0.05 are bolded for emphasis.

**Table S4.** Details of Rise Time Comparisons. Rise times (t_66_-t_33_) of responses to various stimuli, with either an unpaired t-test (if the number of comparisons is one) or an ordinary one-way ANOVA with Dunnett’s multiple comparisons test (if the number of comparisons exceeds one).

| Neuron | Stimulus | Genotype | Mean ± SEM | n | # comparisons | p-value | 95% Confidence Interval | Figure |
| --- | --- | --- | --- | --- | --- | --- | --- | --- |
| AIA | 1.15 μM dia | WT | 4.0 ± 0.2 s | 390 | 3 | | | 1I, 6K |
| | 90 μM IAA | | 3.0 ± 0.5 s | 76 | | 0.332 | [-0.52, 2.36] | |
| | *E. coli* OP50-conditioned medium | | 4.6 ± 0.9 s | 37 | | 0.837 | [-2.59, 1.37] | |
| | AWA::Chr | | 4.1 ± 0.3 s | 260 | | 0.993 | [-1.01, 0.83] | |
| AIA | AWA::Chr | WT | 4.1 ± 0.3 s | 260 | 1 | | | S3G |
| | | *unc-18(e234)* | 3.9 ± 0.3 s | 225 | | 0.813 | [-1.00, 0.78] | |

**Table S5.**
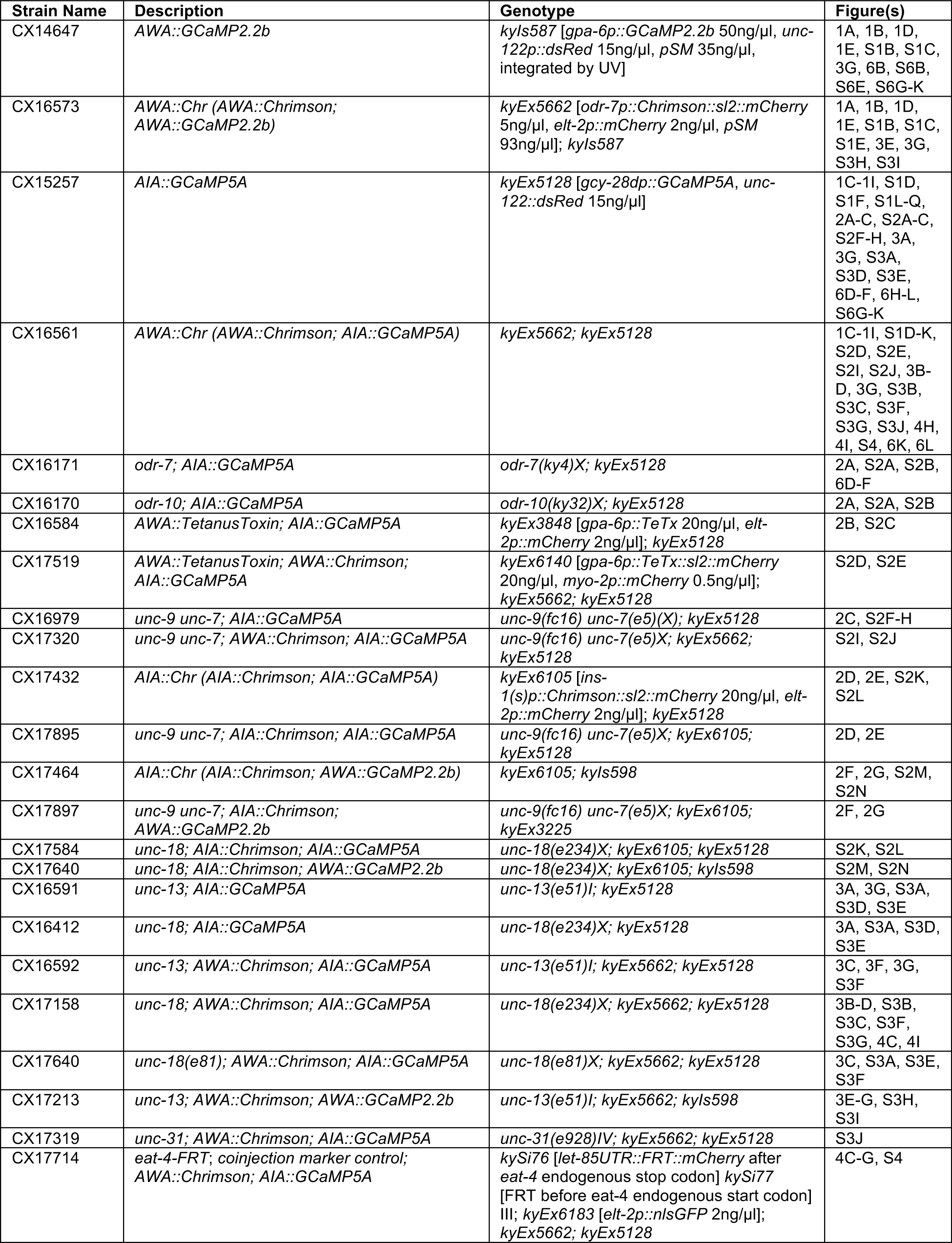
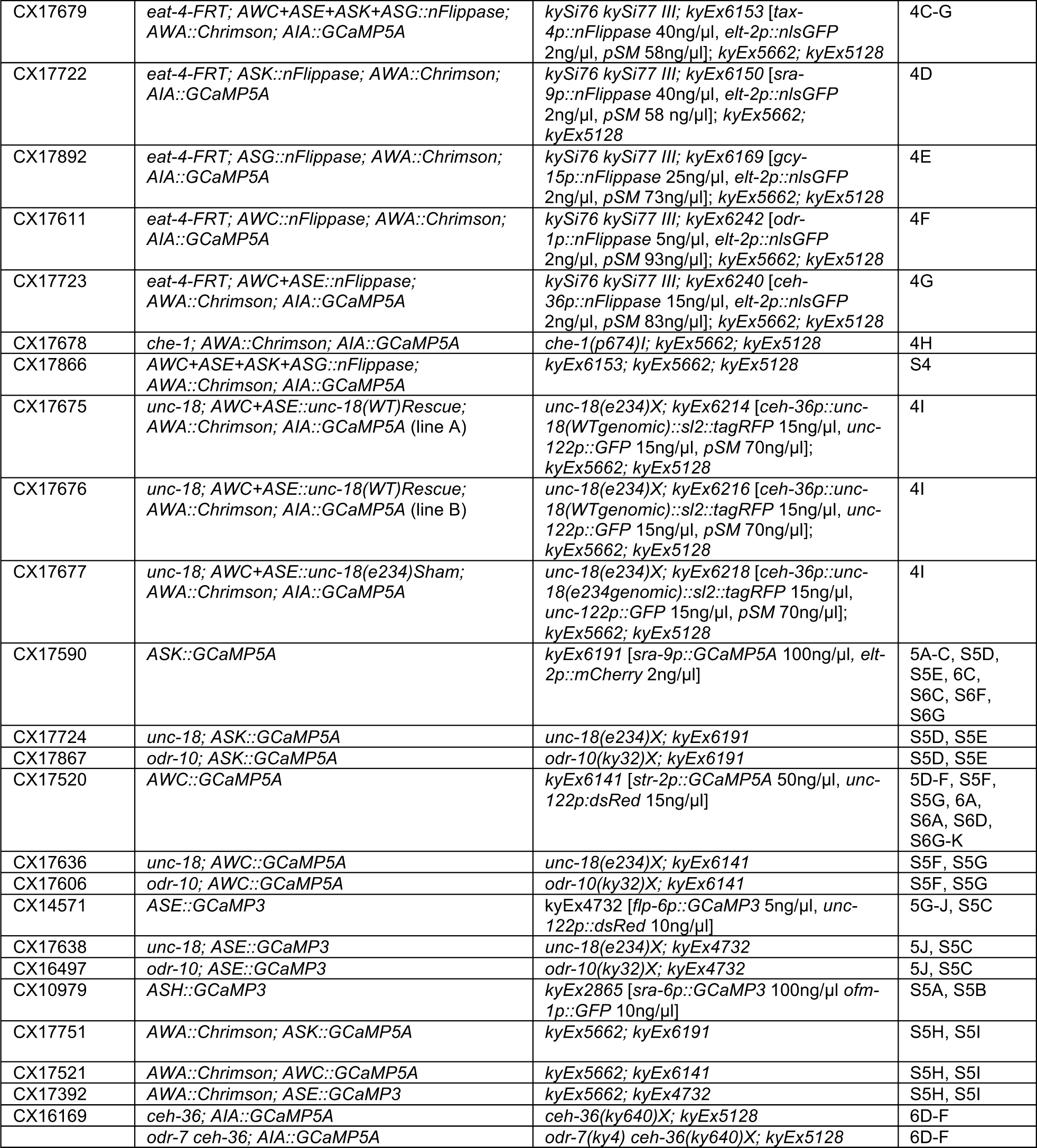
Strain List.

**Figure 1 – figure supplement 1.**
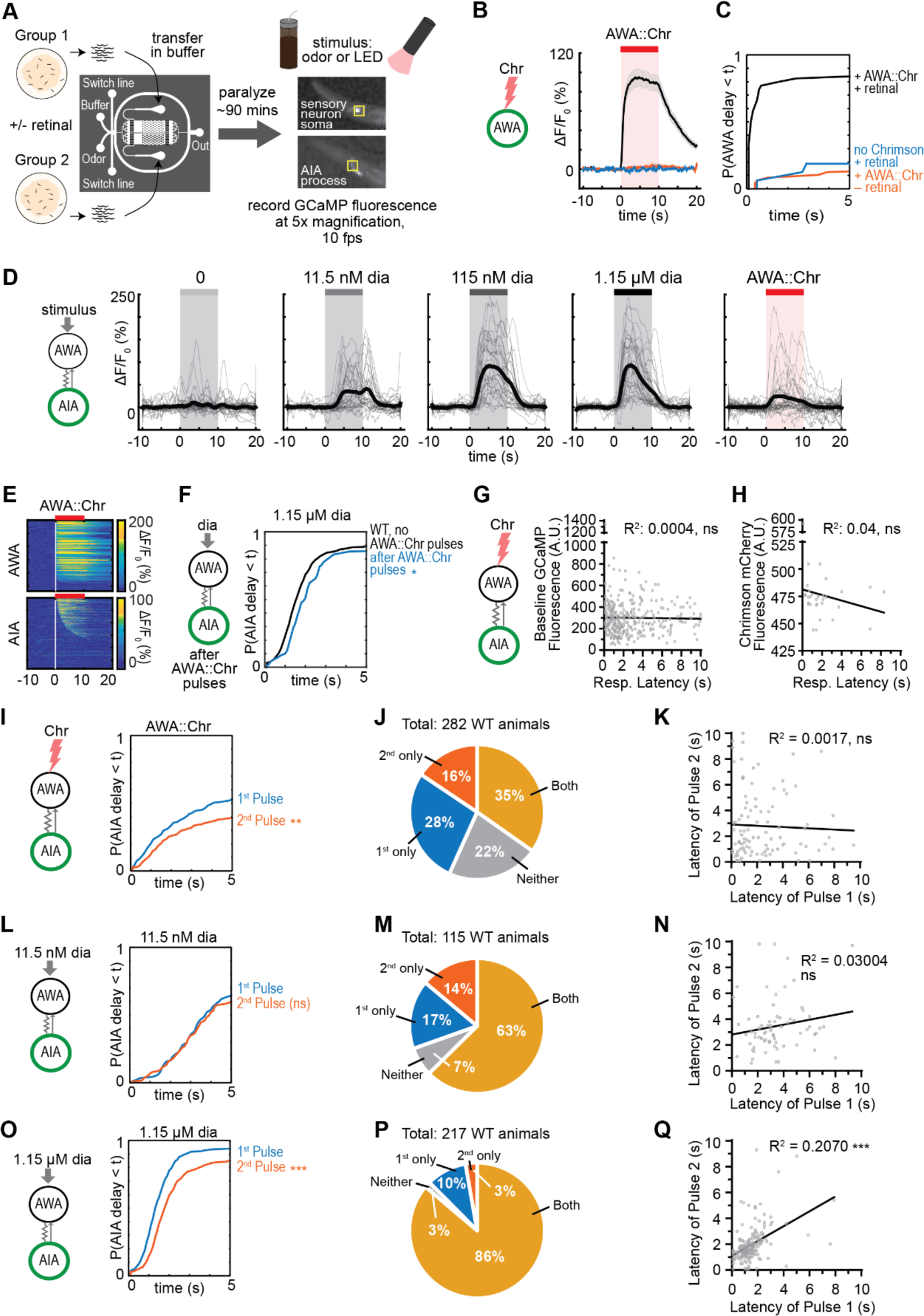
Experimental configuration and control data on reliability of AIA responses. (A) Schematic of experimental configuration. Animals are paralyzed in a microfluidics device and their neural activity is recorded during exposure to odor or light. Two arenas can be recorded simultaneously with up to 10 animals per arena. See Materials and Methods for details. **(B and C)** AWA requires both retinal pre-treatment and expression of the Chrimson transgene for light activation. **(B)** Mean AWA calcium responses; shading indicates ± SEM. Transgene without retinal: n = 48; retinal without transgene: n = 16; transgene with retinal: n = 74. **(C)** Cumulative response time profiles of data from (B), showing first 5 s of light exposure. **(D)** AIA GCaMP5A calcium responses to 10 s pulses of increasing concentrations of diacetyl and to AWA::Chrimson stimulation. Bold lines indicate mean. Responses to AWA::Chrimson stimulation were downsampled to 34 traces at random for visibility from a complete set of 569 traces. **(E)** Heat maps of AWA and AIA responses to AWA::Chrimson stimulation, combined over all experiments. AWA: n = 268; AIA: n = 569. **(F)** Cumulative response time profiles of AIA responses to 1.15 µM diacetyl recorded immediately after recordings of AWA::Chrimson stimulation (blue), representing a subset of animals used in (E). AIA responses to 1.15 µM diacetyl combined over all experiments are shown for comparison (black). **(G)** Response latencies of 318 AIA responses to AWA::Chrimson stimulation do not correlate with GCaMP fluorescence levels at pre-stimulus baseline. **(H)** Response latencies of 31 responses to AWA::Chrimson stimulation do not correlate with Chrimson transgene expression levels. **(I, L, and O)** Cumulative response time profiles of AIA responses to the first or second stimulation with AWA::Chrimson (I), 11.5 nM diacetyl (L), or 1.15 µM diacetyl (O). All other figures and analyses pool responses from both pulses. (J, M, and P) Proportion of animals that respond to only the first, second, both, or neither pulse of stimulation with AWA::Chrimson (J), 11.5 nM diacetyl (M), or 1.15 µM diacetyl (P). **(K)** In 98 animals that responded to both AWA::Chrimson stimulation pulses, there was no correlation between response latencies across pulses. **(N)** In 72 animals that responded to both 11.5 µM diacetyl pulses, there was no correlation between response latencies across pulses. **(Q)** In 187 animals that responded to both 1.15 µM diacetyl pulses, there was a moderate correlation between response latencies across pulses. For (F), (I), (L), and (O), asterisks refer to Kolmogorov-Smirnov test significance over full 10 s stimulus pulse. ns: not significant; *: p<0.05; **: p<0.01; ***: p<0.001. See Table S2 for sample sizes and test details. For (G), (H), (K), (N), and (Q), asterisks refer to significance of linear regression slope differing from 0. ns: not significant; ***: p<0.001.

**Figure 2 – figure supplement 1.**
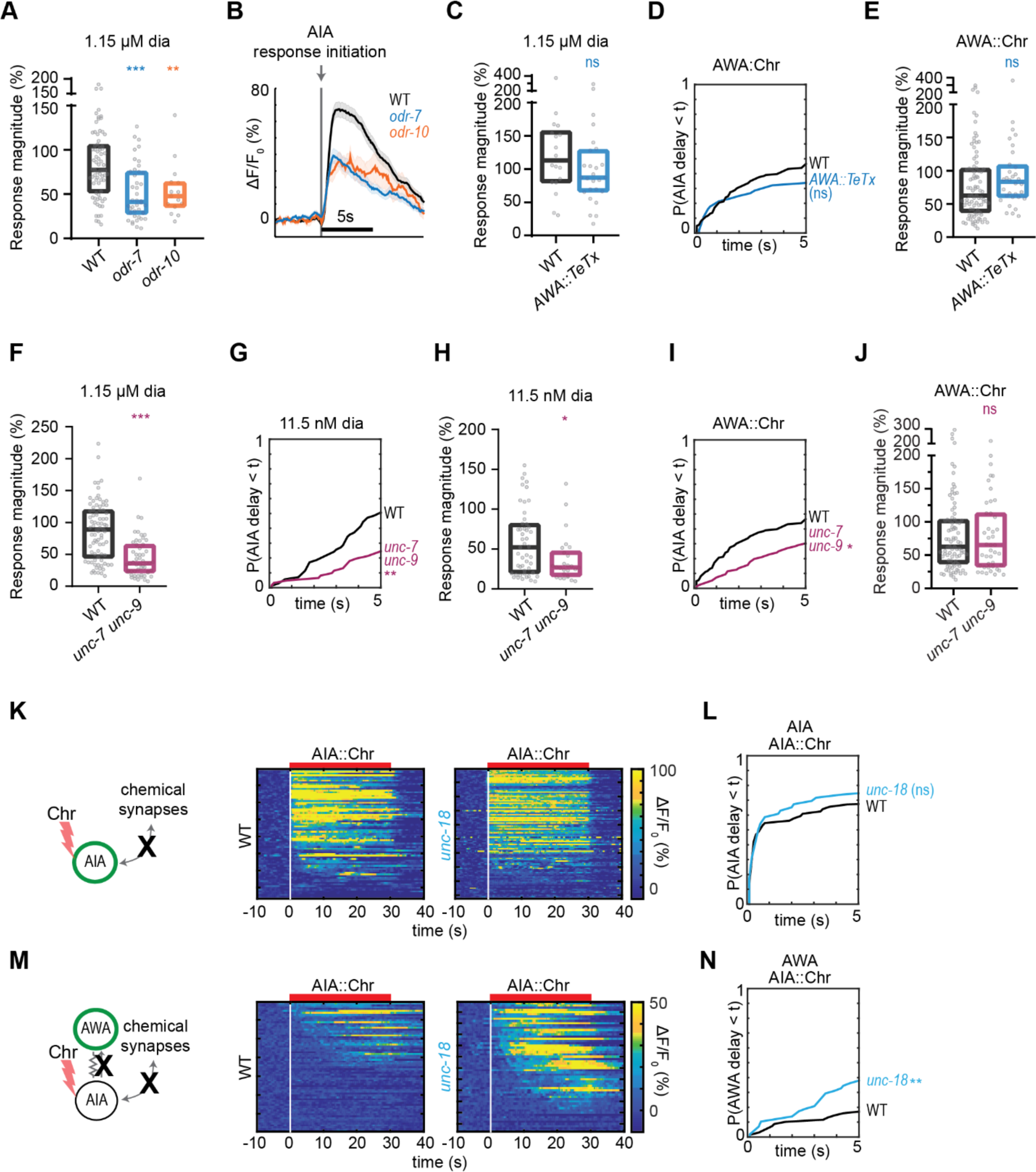
Gap junctions contribute to AWA-to-AIA communication, additional data. (A) Magnitude of AIA responses from Figure 2A, omitting traces that did not produce a detectable response. Boxes show median and interquartile range. **(B)** Mean AIA responses from Figure 2A that initiated within 5 s of stimulus, aligned to the frame at which activation began. Shading indicates ± SEM. **(C)** Magnitude of AIA responses from Figure 2B, omitting traces that did not produce a detectable response. Boxes show median and interquartile range. **(D)** Cumulative response time profiles of AIA responses to AWA::Chrimson stimulation in WT versus animals expressing Tetanus Toxin Light Chain A (TeTx) in AWA. **(E)** Magnitude of AIA responses from (D), omitting traces that did not produce a detectable response. Boxes show median and interquartile range. **(F)** Magnitude of AIA responses from Figure 2C, omitting traces that did not produce a detectable response. Boxes show median and interquartile range. **(G and I)** Cumulative response time profiles of AIA responses to 11.5 nM diacetyl (G) and AWA::Chrimson stimulation (I) in WT versus *unc-7 unc-9* animals (innexin mutants). **(H and J)** Magnitude of AIA responses from (G) and (I), respectively. Boxes show median and interquartile range. (K and M) AIA (K) and AWA (M) responses to 10 s pulses of AIA::Chrimson stimulation in WT and *unc-18* animals (synaptic transmission mutants); one row per calcium trace. **(L)** Cumulative response time profiles of AIA responses shown in (K). (N) Cumulative response time profiles of AWA responses shown in (M). For (A), (C), (E), (F), (H), and (J), asterisks refer to statistical significance of one-way ANOVA with Dunnett’s multiple comparisons test. ns: not significant; *: p<0.05; **: p<0.01; ***: p<0.001. See Table 2 for sample sizes and test details. For (D), (G), (I), (L), and (N), asterisks refer to Kolmogorov-Smirnov test significance versus WT over full 10 s stimulus pulse. ns: not significant; *: p<0.05; **: p<0.01. See Table S2 for sample sizes and test details.

**Figure 3 – figure supplement 1.**
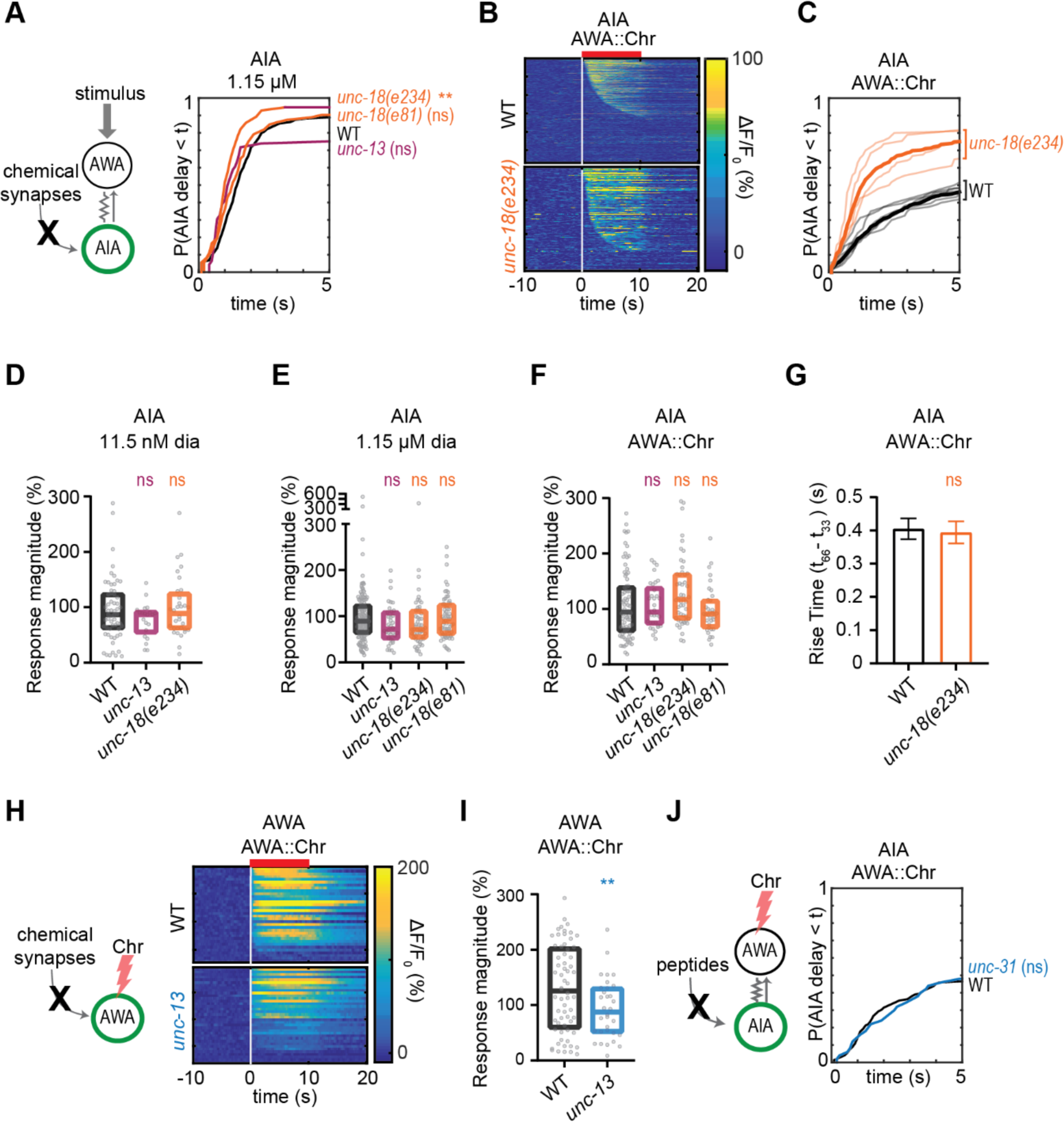
Chemical synapses inhibit AIA, additional data. **(A)** Cumulative response time profiles of AIA to 1.15 µM diacetyl in WT versus *unc-13(e51)*, *unc-18(e234)*, and *unc-18(e81)* animals (synaptic transmission mutants). **(B)** Heat maps of AIA responses to AWA::Chrimson stimulation in WT and *unc-18(e234)* animals, combined over all experiments. N2: n = 569; *unc-18(e234)*: n = 335. **(C)** Cumulative response time profiles of AIA responses shown in (B). Thick lines represent distribution of all experiments, faint lines represent distributions from individual experimental blocks. **(D – F)** Magnitude of AIA responses to 11.5 nM diacetyl (C), 1.15 µM diacetyl (D), and AWA::Chrimson stimulation (E), in WT versus *unc-13*, *unc-18(e234)*, and *unc-18(e81)* animals (synaptic transmission mutants), omitting traces that did not produce a detectable response. Boxes show median and interquartile range. **(G)** Rise time of responses from Figure 3D. Bars indicate SEM. **(H)** Heat maps of AWA responses to AWA::Chrimson stimulation in WT and *unc-13* animals from Figure 3E. **(I)** Magnitude of AWA responses shown in (H), omitting traces that did not produce a detectable response. Boxes show median and interquartile range. (J) Cumulative response time profiles of AIA responses to AWA::Chrimson stimulation in WT versus *unc-31* animals (dense core vesicle exocytosis mutants). For (A) and (J), asterisks refer to Kolmogorov-Smirnov test significance versus WT over full 10 s stimulus pulse. ns: not significant; **: p<0.01. See Table S2 for sample sizes and test details. For (D-F), ns refers to the lack of statistical significance of a one-way ANOVA with Dunnett’s multiple comparisons test. For (G) and (I), asterisks refer to statistical significance of an unpaired t-test. ns: not significant; **: p<0.01. See Table S3 for sample sizes and test details of (D-F), and (I); see Table S4 for samples sizes and test details of (G).

**Figure 4 – figure supplement 1.**
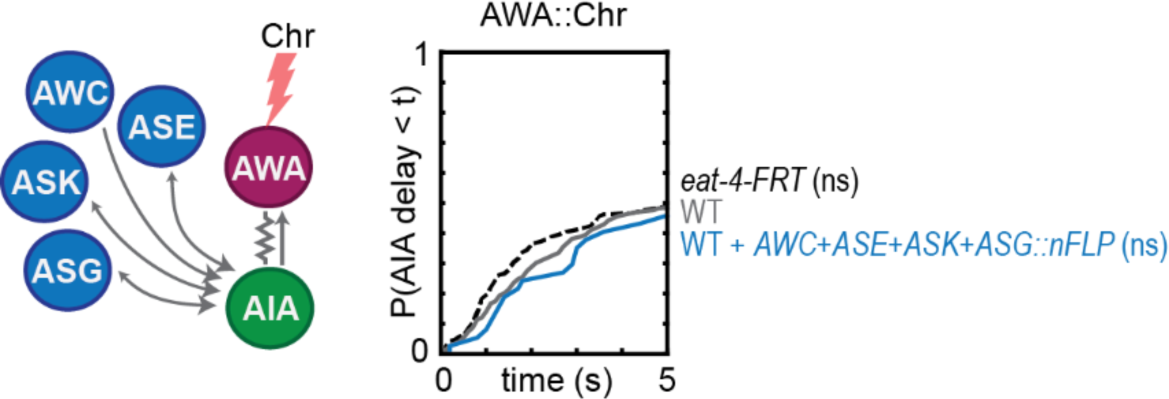
Controls for FRT-FLP recombination. Cumulative response time profiles of AIA responses to AWA::Chrimson stimulation in wild type animals, animals with the genetically modified *eat-4-FRT* locus alone, or animals with the *AWC+ASE+ASK+AWG::nFlippase* transgene alone. All genotypes include the AWA::Chrimson transgene. ns refers to lack of Kolmogorov-Smirnov test significance versus WT over full 10 s stimulus pulse. See Table S2 for sample sizes and test details.

**Figure 5 – figure supplement 1.**
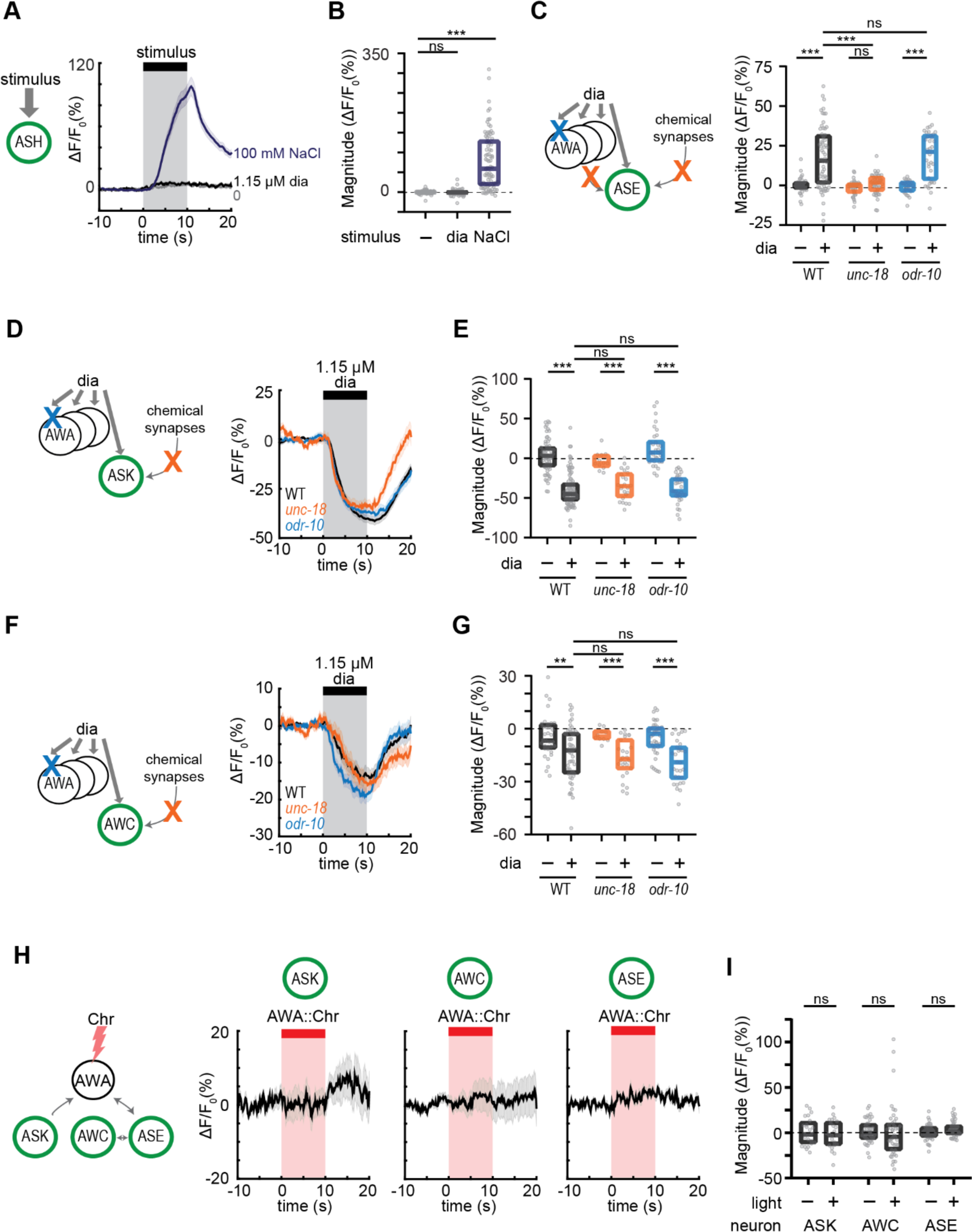
Controls for diacetyl activation of sensory neurons. **(A)** Mean ASH responses to pulses of buffer (0), 1.15 µM diacetyl, or 100 mM NaCl in WT animals. Shading indicates ± SEM. **(B)** Magnitude of individual responses in (A). Boxes show median and interquartile range. **(C)** Schematic of experiment and magnitudes of individual ASE responses in Figure 5J. Boxes show median and interquartile range. **(D and F)** Schematic of experiment and mean ASK (D) and AWC (F) responses to pulses of 1.15 µM diacetyl in WT versus *unc-18* and *odr-10* animals. Shading indicates ± SEM. (E and G) Magnitude of individual responses in (D) and (F), respectively. Boxes show median and interquartile range. **(H)** Mean ASK, AWC, and ASE responses to pulses of AWA::Chrimson stimulation in WT animals. Shading indicates ± SEM. (I) Magnitude of individual responses in (H). Boxes show median and interquartile range. For (B), (C), (E), and (G), asterisks refer to statistical significance of a one-way ANOVA with Dunnett’s multiple comparisons test versus WT. ns: not significant; **: p<0.01; ***: p<0.001. For (I), ns refers to lack of statistical significance in paired t-tests comparing pre-light to within-light periods in same neurons. See Table S3 for sample sizes and test details.

**Figure 6 – figure supplement 1.**
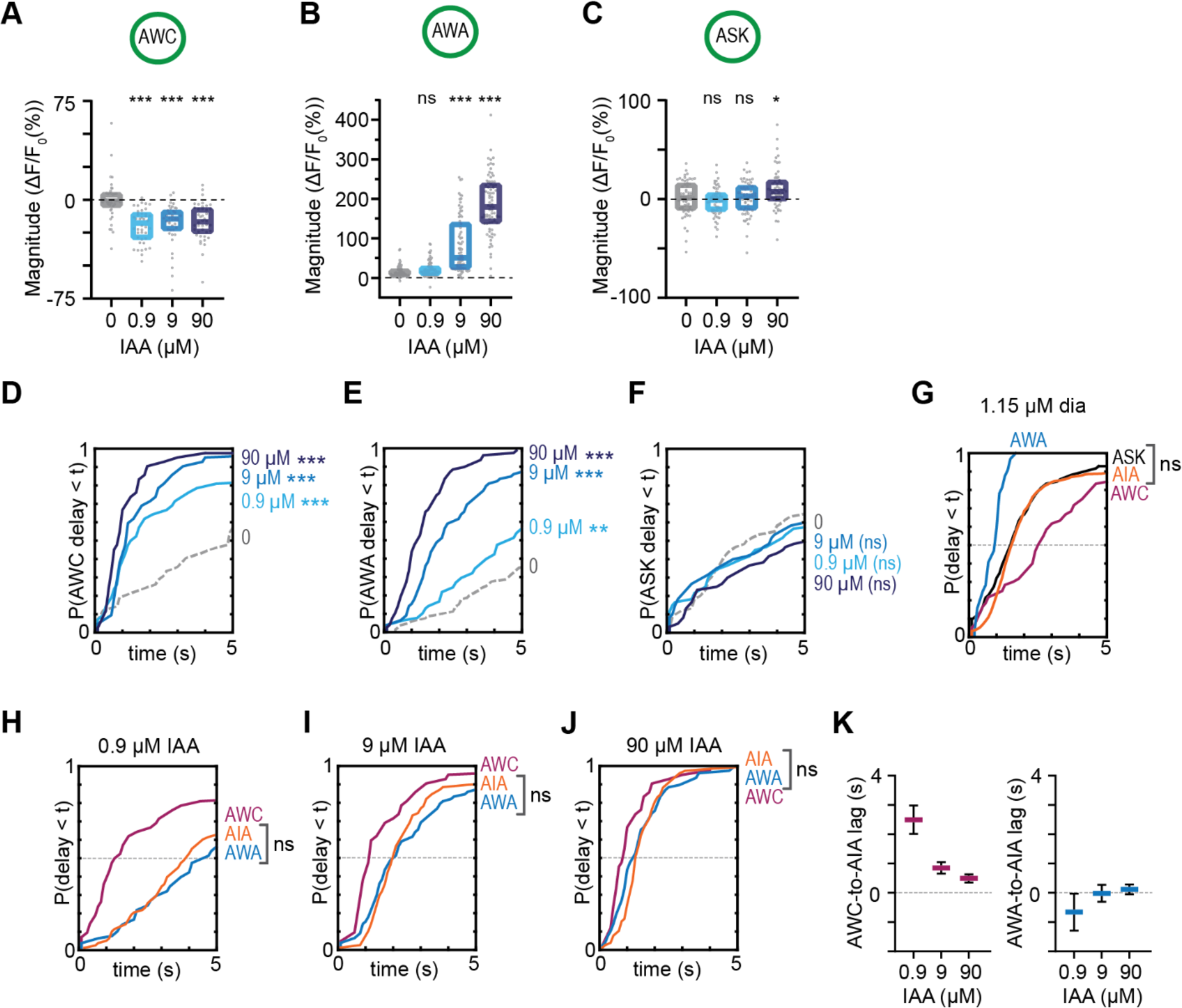
Combinatorial activation of AIA by isoamyl alcohol-sensing neurons, additional data. **(A – C)** Magnitude of individual AWC (A), AWA (B), and ASK (C) responses to isoamyl alcohol shown in Figures 6D-6F. Boxes show median and interquartile range. **(D – F)** Cumulative response time profiles of AWC (D), AWA (E), and ASK (F) responses to isoamyl alcohol shown in Figures 6D-6F. **(G)** Cumulative response time profiles of AWC, AWA, and AIA responses to 1.15 µM diacetyl in WT animals. Note the similarity of ASK and AIA profiles. **(H – J)** Cumulative response time profiles of AWC, AWA, and AIA responses to 0.9 µM (H), 9 µM (I), and 90 µM (J) isoamyl alcohol in WT animals. Note the similarity of AWA and AIA profiles at all concentrations. **(K)** Delay between the time at which 50% of AWC versus 50% of AIA neurons (left) or AWA versus AIA neurons (right) respond to different concentrations of isoamyl alcohol. Bars are mean ± SEM. For (A-C), asterisks refer to statistical significance of a one-way ANOVA with Dunnett’s multiple comparisons test versus buffer. ns: not significant; *: p<0.05; ***: p<0.001. See Table S3 for sample sizes and test details. For (D-J), asterisks refer to statistical significance of a Kolmogorov-Smirnov test versus buffer (D-F), between ASK and AIA (G), or between AWA and AIA (H-J), over full 10 s stimulus pulse. ns: not significant; *: p<0.05; **: p<0.01; ***: p<0.001. See Table S2 for sample sizes and test details.

## REFERENCES

1. Allen MJ, Murphey RK. 2007. The chemical component of the mixed GF-TTMn synapse in *Drosophila melanogaster* uses acetylcholine as its neurotransmitter. European Journal of Neuroscience 26(2):439–445. doi: 10.1111/j.1460-9568.2007.05686.x.

2. Bargmann CI. 2006. Chemosensation in *C. elegans*. *WormBook*, ed. The *C. elegans* Research Community, WormBook. doi: 10.1895/wormbook.1.123.1, http://www.wormbook.org.

3. Bargmann CI, Hartwieg E, Horvitz, HR. 1993. Odorant-selective genes and neurons mediate olfaction in *C. elegans*. Cell 74(3):515–527. doi: 10.1016/0092-8674(93)80053-h.

4. Bhattacharya A, Aghayeva U, Berghoff EG, Hobert O. 2019. Plasticity of the Electrical Connectome of *C. elegans*. Cell 176(5):1174–1189. doi: 10.1016/j.cell.2018.12.024.

5. Brenner S. 1974. The genetics of *Caenorhabditis elegans*. Genetics 77(1):71–94.

6. Buchler NE, Gerland U, Hwa T. 2003. On schemes of combinatorial transcription logic. Proceedings of the National Academy of Sciences of the United States of America 100(9):5136–5141. doi: 10.1073/pnas.0930314100.

7. Cao J, Packer JS, Ramani V, Cusanovich DA, Huynh C, Daza R, Qiu X, Lee C, Furlan SN, Steemers FJ, Adey A, Waterston RH, Trapnell C, Shendure J. 2017. Comprehensive single-cell transcriptional profiling of a multicellular organism. Science 357(6352):661–667. doi: 10.1126/science.aam8940.

8. Chalasani SH, Chronis N, Tsunozaki M, Gray JM, Ramot D, Goodman MB, Bargmann CI. 2007. Dissecting a circuit for olfactory behavior in *Caenorhabditis elegans*. Nature 450(7166):63–70. doi: 10.1038/nature06292.

9. Chalasani SH, Kato S, Albrecht DR, Nakagawa T, Abbott LF, Bargmann CI. 2010. Neuropeptide feedback modifies odor-evoked dynamics in *Caenorhabditis elegans* olfactory neurons. Nature Neuroscience 13(5):615–621. doi: 10.1038/nn.2526.

10. Chen BL, Hall DH, Chklovskii DB. 2006. Wiring optimization can relate neuronal structure and function. Proceedings of the National Academy of Sciences of the United States of America 103(12):4723–4728. doi: 10.1073/pnas.0506806103.

11. Cho CE, Brueggemann C, L’Etoile ND, Bargmann CI. 2016. Parallel encoding of sensory history and behavioral preference during *Caenorhabditis elegans* olfactory learning. eLife 5:e14000. doi: 10.7554/eLife.14000.

12. Chou JH, Bargmann CI, Sengupta P. 2001. The *Caenorhabditis elegans odr-2* Gene Encodes a Novel Ly-6-Related Protein Required for Olfaction. Genetics 157(1):211–224.

13. Cook SJ, Jarrell TA, Brittin CA, Wang Y, Bloniarz AE, Yakovlev MA, Nguyen KCQ, Tang LT, Bayer EA, Duerr JS, Bülow HE, Hobert O, Hall DH, Emmons SW. 2019. Whole-animal connectomes of both *Caenorhabditis elegans* sexes. Nature 571(7763):63–71. doi: 10.1038/s41586-019-1352-7.

14. Dayan P, Abbott LF. 2001. Theoretical Neuroscience: Computation and Mathematical Modeling of Neural Systems. Cambridge, Massachusetts: The MIT Press.

15. Denève S, Machens CK. 2016. Efficient codes and balanced networks. Nature Neuroscience 19(3):375–382. doi: 10.1038/nn.4243.

16. Goodenough DA, Paul DL. 2009. Gap junctions. Cold Spring Harbor Perspectives in Biology 1(1):a002576. doi: 10.1101/cshperspect.a002576.

17. Gordus A, Pokala N, Levy S, Flavell SW, Bargmann CI. 2015. Feedback from Network States Generates Variability in a Probabilistic Olfactory Circuit. Cell 161(2):215–227. doi: 10.1016/j.cell.2015.02.018.

18. Gray JM, Hill JJ, Bargmann CI. 2005. A circuit for navigation in *Caenorhabditis elegans*. Proceedings of the National Academy Sciences of the United States of America 102(9):3184–3191. doi: 10.1073/pnas.0409009101.

19. Hale LA, Lee ES, Pantazis AK, Chronis N, Chalasani SH. 2016. Altered Sensory Code Drives Juvenile-to-Adult Behavioral Maturation in *Caenorhabditis elegans*. eNeuro 3(6):e0175–16.2016. doi: 10.1523/ENEURO.0175-16.2016.

20. Hendricks M, Ha H, Maffey N, Zhang Y. 2012. Compartmentalized calcium dynamics in a *C. elegans* interneuron encode head movement. Nature 487(7405):99–103. doi: 10.1038/nature11081.

21. Iino Y, Yoshida K. 2009. Parallel use of two behavioral mechanisms for chemotaxis in *Caenorhabditis elegans*. Journal of Neuroscience 29(17):5370–5380. doi: 10.1523/JNEUROSCI.3633-08.2009.

22. Kaplan HS, Nichols ALA, Zimmer M. 2018. Sensorimotor integration in *Caenorhabditis elegans*: a reappraisal towards dynamic and distributed computations. Philosophical Transactions of the Royal Society B 373(1758):20170371. doi: 10.1098/rstb.2017.0371.

23. Kato S, Kaplan HS, Schrödel, Skora S, Lindsay TH, Yemini E, Lockery S, Zimmer M. 2015. Global Brain Dynamics Embed the Motor Command Sequence of *Caenorhabditis elegans*. Cell 163(3):656–669. doi: 10.1016/j.cell.2015.09.034.

24. Kato S, Xu Y, Cho CE, Abbott LF, Bargmann CI. 2014. Temporal responses of *C. elegans* chemosensory neurons are preserved in behavioral dynamics. Neuron 81(3):616–628. doi: 10.1016/j.neuron.2013.11.020.

25. Klapoetke NC, Murata Y, Kim SS, Pulver SR, Birdsey-Benson A, Cho YK, Morimoto TK, Chuong AS, Carpenter EJ, Tian Z, Wang J, Xie Y, Yan Z, Zhang Y, Chow BY, Surek B, Melkonian M, Jayaraman V, Constantine-Paton M, Wong GK, Boyden ES. 2014. Independent optical excitation of distinct neural populations. Nature Methods 11(3):338–346. doi: 10.1038/nmeth.2836.

26. Koch C. 1999. Biophysics of Computation: Information Processing in Single Neurons. New York, New York: Oxford University Press.

27. Lanjuin A, VanHoven MK, Bargmann CI, Thompson JK, Sengupta P. 2003. *Otx/otd* Homeobox Genes Specify Distinct Sensory Neuron Identities in *C. elegans*. Developmental Cell 5(4):621–633. doi: 10.1016/S1534-5807(03)00293-4.

28. Larsch J, Flavell SW, Liu Q, Gordus A, Albrecht DR, Bargmann CI. 2015. A Circuit for Gradient Climbing in *C. elegans*. Cell Reports 12(11):1748–1760. doi: 10.1016/j.celrep.2015.08.032.

29. Larsch J, Ventimiglia D, Bargmann CI, Albrecht DR. 2013. High-throughput imaging of neuronal activity in *Caenorhabditis elegans*. Proceedings of the National Academy of Sciences of the United States of America 110(45):E4266–4273. doi: 10.1073/pnas.1318325110.

30. Lee RY, Sawin ER, Chalfie M, Horvitz HR, Avery L. 1999. EAT-4, a homolog of a mammalian sodium-dependent inorganic phosphate cotransporter, is necessary for glutamatergic neurotransmission in *Caenorhabditis elegans*. Journal of Neuroscience 19(1):159–167.

31. Li Z, Liu J, Zheng M, Xu XZ. 2014. Encoding of both analog- and digital-like behavioral outputs by one *C. elegans* interneuron. Cell 159(4):751–765. doi: 10.1016/j.cell.2014.09.056.

32. Liu P, Chen B, Altun ZF, Gross MJ, Shan A, Schuman B, Hall DH, Wang ZW. 2013. Six innexins contribute to electrical coupling of *C. elegans* body-wall muscle. PLoS One 8(10):e76877. doi: 10.1371/journal.pone.0076877.

33. Liu P, Chen B, Mailler R, Wang ZW. 2017. Antidromic-rectifying gap junctions amplify chemical transmission at functionally mixed electrical-chemical synapses. Nature Communications 8:14818. doi: 10.1038/ncomms14818.

34. Liu Q, Kidd PB, Dobosiewicz M, Bargmann CI. 2018. *C. elegans* AWA Olfactory Neurons Fire Calcium-Mediated All-or-None Action Potentials. Cell 175(1):57–70. doi: 10.1016/j.cell.2018.08.018.

35. López-Cruz A, Sordillo A, Pokala N, Liu Q, McGrath PT, Bargmann CI. 2019. Parallel Multimodal Circuits Control an Innate Foraging Behavior. Neuron 102(2):407–419. doi: 10.1016/j.neuron.2019.01.053.

36. Macosko EZ, Pokala N, Feinberg EH, Chalasani SH, Butcher RA, Clardy J, Bargmann CI. 2009. A hub-and-spoke circuit drives pheromone attraction and social behaviour in *C. elegans*. Nature 458(7242):1171–1175. doi: 10.1038/nature07886.

37. McCulloch WS, Pitts W. 1943. A logical calculus of the ideas immanent in nervous activity. The Bulletin of Mathematical Biophysics 5(4):115–133.

38. Miller AC, Whitebirch AC, Shah AN, Marsden KC, Granato M, O’Brien J, Moens CB. 2017. A genetic basis for molecular asymmetry at vertebrate electrical synapses. eLife 6:e25364. doi: 10.7554/eLife.25364.

39. Okun M, Lampl I. 2008. Instantaneous correlation of excitation and inhibition during ongoing and sensory-evoked activities. Nature Neuroscience 11(5):535–537. doi: 10.1038/nn.2105.

40. Pereda AE, Curti S, Hoge G, Cachope R, Flores CE, Rash JE. 2013. Gap junction-mediated electrical transmission: Regulatory mechanisms and plasticity. Biochimica et Biophysica Acta 1828:134–146. doi: 10.1016/j.bbamem.2012.05.026.

41. Pereira L, Kratsios P, Serrano-Saiz E, Sheftel H, Mayo AE, Hall DH, White JG, LeBoeuf B, Garcia LR, Alon U, Hobert O. 2015. A cellular and regulatory map of the cholinergic nervous system of *C. elegans*. eLife 4:e12432. doi: 10.7554/eLife.12432.

42. Phelan P, Goulding LA, Tam JL, Allen MJ, Dawber RJ, Davies JA, Bacon JP. 2008. Molecular mechanism of rectification at identified electrical synapses in the *Drosophila* giant fiber system. Current Biology 18(24):1955–1960. doi: 10.1016/j.cub.2008.10.067.

43. Pi HJ, Hangya B, Kvitsiani D, Sanders JI, Huang ZJ, Kepecs A. 2013. Cortical interneurons that specialize in disinhibitory control. Nature 503(7477):521–524. doi: 10.1038/nature12676.

44. Robertson H, Thomas J. 2006. The putative chemoreceptor families of *C. elegans. Wormbook* ed. The *C. elegans* Research Community, WormBook, doi: 10.1895/wormbook.1.66.1, http://www.wormbook.org.

45. Richmond J. 2007. Synaptic function. *WormBook*, ed. The *C. elegans* Research Community, WormBook, doi:10.1895/wormbook.1.69.1, http://www.wormbook.org.

46. Sengupta P, Chou JH, Bargmann CI. 1996. *odr-10* Encodes a Seven Transmembrane Domain Olfactory Receptor Required for Responses to the Odorant Diacetyl. Cell 84(6):899–909. doi: 10.1016/s0092-8674(00)81068-5.

47. Sengupta P, Colbert HA, Bargmann CI. 1994. The *C. elegans* Gene *odr-7* Encodes an Olfactory-Specific Member of the Nuclear Receptor Superfamily. Cell 79(6):971–980. doi: 10.1016/0092-8674(94)90028-0.

48. Serrano-Saiz E, Poole RJ, Felton T, Zhang F, De La Cruz ED, Hobert O. 2013. Modular control of glutamatergic neuronal identity in *C. elegans* by distinct homeodomain proteins. Cell 155(3):659–673. doi: 10.1016/j.cell.2013.09.052.

49. Shinkai Y, Yamamoto Y, Fujiwara M, Tabata T, Murayama T, Hirotsu T, Ikeda DD, Tsunozaki M, Iino Y, Bargmann CI, Katsura I, Ishihara T. 2011. Behavioral choice between conflicting alternatives is regulated by a receptor guanylyl cyclase, GCY-28, and a receptor tyrosine kinase, SCD-2, in AIA interneurons of *Caenorhabditis elegans*. Journal of Neuroscience 31(8):3007–3015. doi: 10.1523/JNEUROSCI.4691-10.2011.

50. Troemel ER, Kimmel BE, Bargmann CI. 1997. Reprogramming chemotaxis responses: sensory neurons define olfactory preferences in *C. elegans*. Cell 91(2):161–169. doi: 10.1016/s0092-8674(00)80399-2.

51. Tsalik EL, Hobert O. 2003. Functional mapping of neurons that control locomotory behavior in *Caenorhabditis elegans*. Journal of Neurobiology 56(2):178–197. doi: 10.1002/neu.10245.

52. Turi GF, Li WK, Chavlis S, Pandi I, O’Hare J, Priestley JB, Grosmark AD, Liao Z, Ladow M, Zhang JF, Zemelman BV, Poirazi P, Losonczy A. 2019. Vasoactive Intestinal Polypeptide-Expressing Interneurons in the Hippocampus Support Goal-Oriented Spatial Learning. Neuron 101(6):1150–1165. doi: 10.1016/j.neuron.2019.01.009.

53. Wakabayashi T, Kimura Y, Ohba Y, Adachi R, Satoh Y, Shingai R. 2009. *In vivo* calcium imaging of OFF-responding ASK chemosensory neurons in *C. elegans*. Biochimica et Biophysica Acta 1790(8):765–769. doi: 10.1016/j.bbagen.2009.03.032.

54. White JG, Southgate E, Thomson J, Brenner S. 1986. The structure of the nervous system of the nematode *Caenorhabditis elegans*. Philosophical Transactions of the Royal Society B 314(1165):1–340. doi: 10.1098/rstb.1986.0056.

55. Worthy SE, Haynes L, Chambers M, Bethune D, Kan E, Chung K, Ota R, Taylor CJ, Glater EE. 2018. Identification of attractive odorants released by preferred bacterial food found in the natural habitats of *C. elegans*. PLoS One 13(7):e0201158. doi: 10.1371/journal.pone.0201158.

56. Yoshida K, Hirotsu T, Tagawa T, Oda S, Wakabayashi T, Iino Y, Ishihara T. 2012. Odour concentration-dependent olfactory preference change in *C. elegans*. Nature Communications 3:739. doi: 10.1038/ncomms1750.

57. Zaslaver A, Liani I, Shtangel O, Ginzburg S, Yee L, Sternberg PW. 2015. Hierarchical sparse coding in the sensory system of *Caenorhabditis elegans*. Proceedings of the National Academy of Sciences of the United States of America 112(4):1185–1189. doi: 10.1073/pnas.1423656112.

